# Population history of the Sardinian people inferred from whole-genome sequencing

**DOI:** 10.1101/092148

**Authors:** Charleston W K Chiang, Joseph H Marcus, Carlo Sidore, Hussein Al-Asadi, Magdalena Zoledziewska, Maristella Pitzalis, Fabio Busonero, Andrea Maschio, Giorgio Pistis, Maristella Steri, Andrea Angius, Kirk E Lohmueller, Goncalo R Abecasis, David Schlessinger, Francesco Cucca, John Novembre

## Abstract

The population of the Mediterranean island of Sardinia has made important contributions to genome-wide association studies of traits and diseases. The history of the Sardinian population has also been the focus of much research, and in recent ancient DNA (aDNA) studies, Sardinia has provided unique insight into the peopling of Europe and the spread of agriculture. In this study, we analyze whole-genome sequences of 3,514 Sardinians to address hypotheses regarding the founding of Sardinia and its relation to the peopling of Europe, including examining fine-scale substructure, population size history, and signals of admixture. We find the population of the mountainous Gennargentu region shows elevated genetic isolation with higher levels of ancestry associated with mainland Neolithic farmers and depleted ancestry associated with more recent Bronze Age Steppe migrations on the mainland. Notably, the Gennargentu region also has elevated levels of pre-Neolithic hunter-gatherer ancestry and increased affinity to Basque populations. Further, allele sharing with pre-Neolithic and Neolithic mainland populations is larger on the X chromosome compared to the autosome, providing evidence for a sex-biased demographic history in Sardinia. These results give new insight to the demography of ancestral Sardinians and help further the understanding of sharing of disease risk alleles between Sardinia and mainland populations.

## INTRODUCTION

How complex traits change through time is a central question in evolutionary biology and genetics. Human genetics provides a compelling context for studying this process; however, indepth studies of complex trait evolution in humans requires populations where it is possible to integrate trait mapping with a detailed knowledge of population history. Among human populations, the people of the Mediterranean island of Sardinia are particularly well suited for genetic studies as evident from a number of successes in complex trait and disease mapping (Lettre and Hirschhorn 2015). For example, early studies illuminated the genetic basis of thalassemia and more recent studies have mapped novel quantitative trait loci (QTL) for traits such as hemoglobin levels (Sidore et al. 2015), inflammation levels (Naitza et al. 2012, Sidore et al. 2015) and height (Zoledziewska et al. 2015). Several of these complex traits and diseases show unique incidences in Sardinia (e.g. Marrosu et al. 2004, Pugliatti et al. 2006, Cao and Galanello 2010). Understanding how and why these traits reached their frequencies in Sardinia would provide useful case studies of the dynamics of complex trait evolution. Yet to empower such studies, a detailed background population history of Sardinia is needed.

A clear feature of the history of Sardinia is its differentiation from mainland populations, as evidenced by a distinctive cultural, linguistic, and archaeological legacy (Dyson and Rowland 2007). Early genetic studies made clear that Sardinia has also been a genetically isolated population on the basis of classical autosomal markers, uniparental markers, and elevated linkage disequilibrium (Barbujani and Sokal 1990, Cavalli-Sforza and Piazza 1993, Eaves et al. 2000, Zavattari et al. 2000, Calo et al. 2008). Partly on this basis, Sardinia was included in the Human Genome Diversity Panel (HGDP; Cann 1998), which has been used as a reference sample in many studies, including recent ancient DNA (“aDNA”) studies of Europe (Li et al. 2008, Keller et al. 2012, Skoglund et al. 2012, Lazaridis et al. 2014, Haak et al. 2015). Despite substantial research and interest in Sardinia, genetic studies of its demographic history are still incomplete.

One of the most remarkable findings to date regarding Sardinia’s demographic history is that it has the highest detected levels of genetic similarity to ancient Neolithic farming peoples of Europe (Keller et al. 2012, Skoglund et al. 2012, Lazaridis et al. 2014, Sikora et al. 2014, Allentoft et al. 2015, Haak et al. 2015, Mathieson et al. 2015, Hofmanova et al. 2016). This affiliation of Sardinians with the early Neolithic farmers is currently interpreted in a model with three ancestral populations that contribute ancestry to modern European populations (Lazaridis et al. 2014, Allentoft et al. 2015, Haak et al. 2015, Mathieson et al. 2015). This model postulates that early Neolithic farmers (“EF”) from the Near East and Anatolia expanded into Europe ~7,500 to ~8,000 years ago and mixed in varying proportions with the existing hunter-gatherer peoples (“HG”) in Europe. Then a substantial expansion of Steppe pastoralists (“SP”, associated with the Yamnaya culture) in the Bronze Age ~4,500 to ~5,000 years ago introduced a third major component of ancestry across Europe. This model has been useful in explaining numerous patterns observed in ancient and modern DNA data throughout Europe; here we look closely at how well it explains Sardinian population history.

In this model, Sardinia is effectively colonized by the EF during the European Neolithic, with minor contributions from pre-Neolithic HG groups. Sardinia then remained largely isolated from subsequent migrations on the continent (Sikora et al. 2014), including the Bronze Age expansions of the SP (Lazaridis et al. 2014, Allentoft et al. 2015, Haak et al. 2015). Support for this model is based on analysis of autosomal single-nucleotide polymorphism (SNP) array data, and has additional support from a mtDNA-based ancient DNA study in Sardinia which showed evidence of relative isolation from mainland Europe since the Bronze Age, particularly for the Ogliastra region (Ghirotto et al. 2010). There is also support for this model in the relatively low frequency in Sardinia of U haplogroups that are markers of hunter-gather ancestry (Morelli et al. 2000, Fraumene et al. 2003, Pala et al. 2009, Haak et al. 2015, Olivieri et al. 2017). The archaeological record in Sardinia is also broadly supportive of such a model, in that there are few notable sites from the pre-Neolithic, followed by an expansion of sites in the Neolithic and development of a unique local cultural assemblage (Nuragic culture) by the Bronze Age in Sardinia (see Caramelli et al. 2007, Francalacci et al. 2013).

The conclusion that Sardinia was effectively descended from early Neolithic farmers is not without question though. The first human remains on Sardinia date to the Upper Paleolithic time and flint-stone instruments are found in the Lower Paleolithic period (Vona 1997, Calo et al. 2008), so the potential earliest residents arrived in Sardinia ~14,000-18,000 years ago, much earlier than the Neolithic period. Studies using Y-chromosome haplotypes have found Mesolithic or Paleolithic dates for the common ancestor of Sardinian-specific haplotypes and have interpreted these results as evidence for a strong pre-Neolithic component of Sardinian ancestry (Semino et al. 2000, Rootsi et al. 2004, Calo et al. 2008, Contu et al. 2008, Morelli et al. 2010).

Of particular relevance are the Y-chromosome haplogroups I2a1a (also known as I-M26) and R1b1a2 (also known as R-M269), which have both been used to argue for a large component of HG ancestry in Sardinia. Specifically, I2a1a, is: 1) prevalent in Sardinia (~39%) but rare elsewhere in Europe (Semino et al. 2000, Rootsi et al. 2004, Contu et al. 2008, Francalacci et al. 2013), 2) dated to be pre-Neolithic in origin (Semino et al. 2000, Rootsi et al. 2004, Contu et al. 2008), and 3) found predominately among pre-Neolithic European HG groups in aDNA (Haak et al. 2015, Mathieson et al. 2015). Thus, it is unclear why this haplotype has such a high frequency in Sardinia if most of Sardinian ancestry derives from the EF. Moreover, the relatively high prevalence of R haplogroup R1b1a2 (R-M269) haplogroup in Sardinia (~18%) has been interpreted to reflect a large component of pre-Neolithic HG ancestry in Sardinia (Contu et al. 2008, Morelli et al. 2010), although recent aDNA studies suggest the earliest observation of the haplogroup may be in ancient Yamnaya individuals (Haak et al. 2015).

Whether the high frequencies of I2a1a and R1b1a2 haplogroups in Sardinia are indicative of a pre-Neolithic ancestry of Sardinia is an important open question. Several authors have argued against interpreting the Y chromosome as evidence of Paleolithic ancestry (Passarino et al. 2001, Chikhi et al. 2002, Fraumene et al. 2003, Caramelli et al. 2007, Ghirotto et al. 2010). Also, discrepancies between Y chromosome and autosomal SNPs can arise from accelerated genetic drift on the Y chromosome, sex-biased migrations, or natural selection.

Here, to provide novel insight to the peopling of Sardinia and its relationship to mainland populations, we use an extensive dataset of 3,514 individual whole-genome sequences sampled as part of the SardiNIA project (Sidore et al. 2015). The advent of whole-genome sequencing and large genome-wide SNP marker datasets has revolutionized demographic studies in population genetics (Schraiber and Akey 2015). We leverage these tools to address whether the isolation of Sardinia is consistent with a dominantly EF or HG peopling and whether there is evidence for sex-biased processes that may explain the apparent discrepancy in signals between autosomal and Y chromosome results. As part of our analysis we address the hypothesis of an ancestral connection between Sardinia and the Basque populations of Spain and France (the Basque also show high affinity with early Neolithic farmer aDNA (Lazaridis et al. 2014, Gunther et al. 2015)). We also address whether gene flow from North African populations to Sardinia (Moorjani et al. 2011, Francalacci et al. 2013, Loh et al. 2013, Hellenthal et al. 2014) has been substantial as Sardinia may be the result of a recent multiway admixture involving sources from around the Mediterranean (Hellenthal et al. 2014).

A key factor in our analysis is the consideration of internal sub-structure within Sardinia. Numerous studies have noted the relative heterogeneity of sub-populations within Sardinian (Barbujani et al. 1995, Morelli et al. 2000, Fraumene et al. 2003, Caramelli et al. 2007, Pistis et al. 2009, Ghirotto et al. 2010). Our dataset includes broadly dispersed samples from across Sardinia (Zoledziewska et al. 2009, Sanna et al. 2010), as well as a deep sampling of several villages from the Lanusei Valley region of the Ogliastra province (Pilia et al. 2006, Sidore et al. 2015) (**Figure 1**). We show samples from the Ogliastra province and the broader, mountainous Gennargentu region have signs of elevated isolation. Thus, we frame our analyses of the demographic history of Sardinia by contrasting results for sampled individuals from the Gennargentu region with those outside of the region.

**Figure 1:**
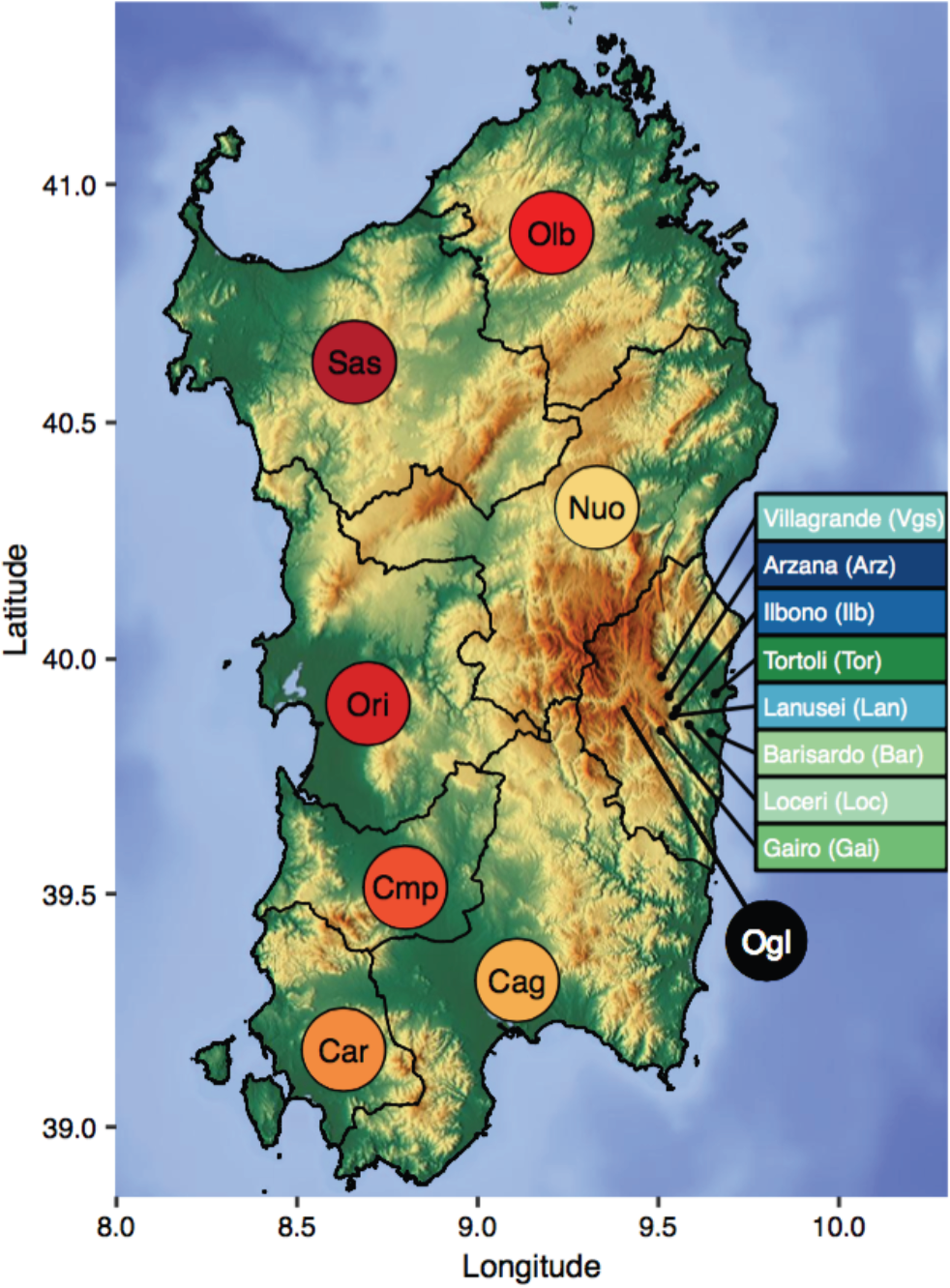
Geographical map of Sardinia. The provincial boundaries are given as black lines. The provinces are abbreviated as Cag (Cagliari), Cmp (Campidano), Car (Carbonia), Ori (Oristano), Sas (Sassari), Olb (Olbia-tempio), Nuo (Nuoro), and Ogl (Ogliastra). For sampled villages within Ogliastra, the names and abbreviations are indicated in colored boxes. Color corresponds to the color used in the PCA plot (Figure 2). The Gennargentu region referred to in the main text is the mountainous area shown in brown that is centered in western Ogliastra (Ogl) and southeastern Nuoro (Nuo).

## RESULTS

Before addressing broader-scale questions regarding Sardinian demographic history, we first examine population structure within Sardinia. We focus on a subset (N=1,577) of unrelated individuals with at least three grandparents originating from the same geographical location to lessen the confounding of internal migration events in the last century.

We find the strongest axis of genetic variation is between individuals from Ogliastra (Ogl) and those outside of Ogliastra (**Figure 2A-B**). Samples from outside Ogliastra show lower levels of differentiation by F_S_t amongst themselves (**Figure 2C**), as well as higher levels of allele sharing (**Figure 3**) despite the greater geographical distance between populations. When we use a spatially explicit statistical method (EEMS, (Petkova et al. 2016)) for visualizing genetic diversity patterns, the resulting effective migration surface (**Figure 2D**) is consistent with high effective migration in western regions of Sardinia connecting the major populations centers of Cagliari (Cag), Oristano (Ori), and Sassari (Sas). Low effective migration rates separate these provinces from a broad area that extends to the mountainous Gennargentu Massif region, including inland Ogliastra to the west. The Gennargentu region is also where some of the Sardinian individuals in the Human Genome Diversity Project (HGDP) originate (A. Piazza, personal communication). We find the HGDP Sardinia individuals partially overlap with our dataset and include a subset that cluster near the Ogliastra sub-population (**Figure S1, S2, Table S1, S2**). Thus, we use the term “Gennargentu-region” to describe the ancestry component (red component in **Figure 2B**). Based on these results, and to simplify analyses going forward, we use individuals from the town of Arzana as a representative of the Gennargentu-region ancestry component and Cagliari as a representative of ancestry outside of the Gennargentu region.

**Figure 2:**
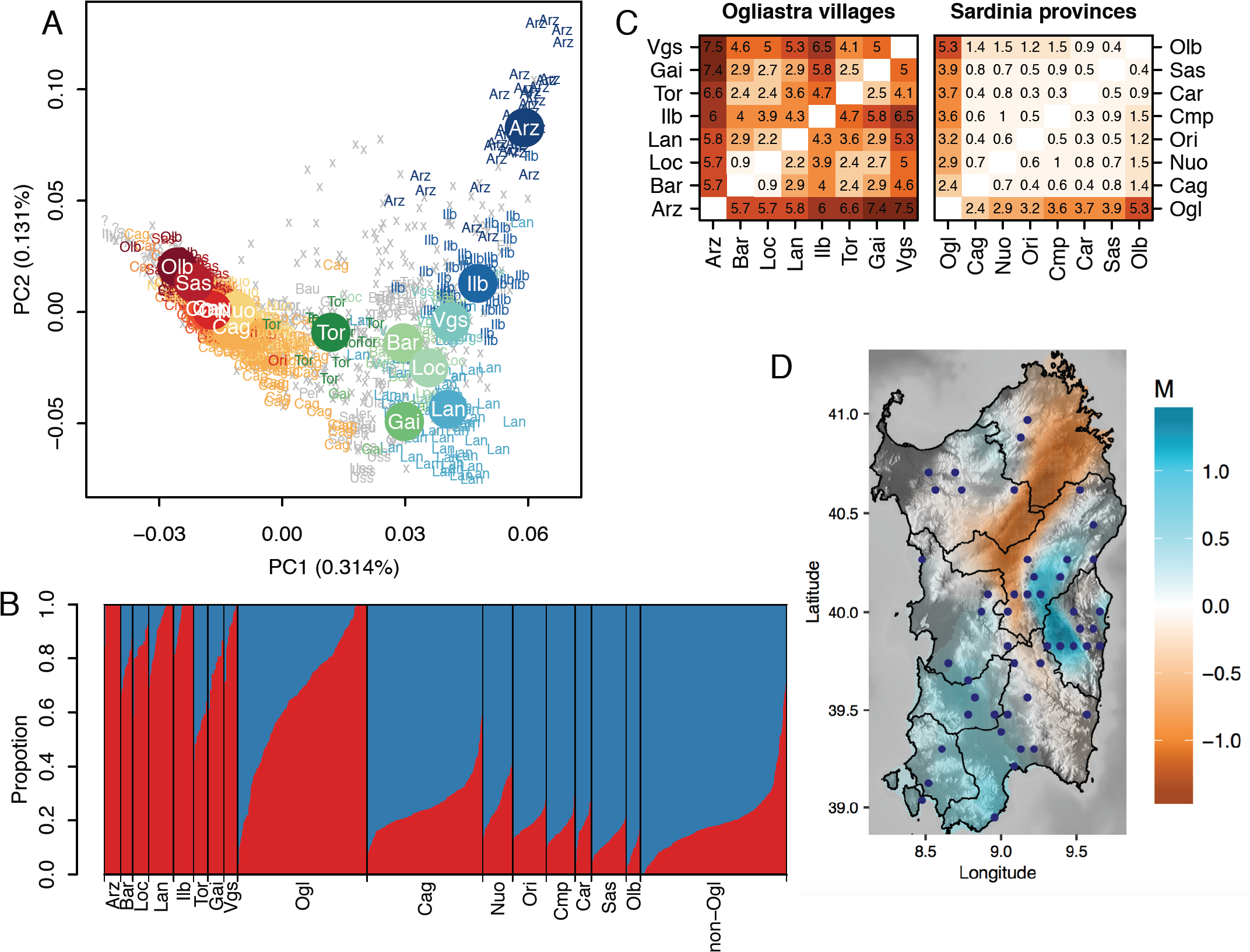
Within-island population structure. (A) Top two components of PCA based on 1,577 unrelated Sardinians. Each individual is labeled with a location if at least 3 of the 4 grandparents were born in the same geographical location; otherwise they are labeled with x. If grandparental ancestry is missing, they are labeled with ?. Subpopulations with at least 8 individuals were colored, otherwise they are displayed in the background in grey color. PC1 differentiates individuals within Ogliastra from those outside of Ogliastra. A notable exception is the sub-population from Tortoli, a recently developed coastal city and the main seaport of the Ogliastra region. Samples from Tortoli show closer affinity in the PCA to samples from the western part of the island. (B) Admixture result at K = 2, which had the lowest cross-validation errors from K = 2 to K = 7 (not shown). Individuals from Ogliastra and outside of Ogliastra that were not assigned to a major location or had mixed grandparental origins are grouped under Ogl and non-Ogl, respectively. Bars for locations with individuals fewer than 40 (Bar, Gai, Loc, Olb, Tor, Vgs) were expanded to a fixed minimum width to aid visualization. (C) Genetic differentiation among Ogliastra villages (left) or among Sardinia provinces (right) as measured by Fst. Ogliastra appears to be the most differentiated from other provinces and within Ogliastra the level of differentiation between villages is substantial (reaching as high as 0.0075 between Villagrande and Arzana), with Arzana being consistently well differentiated from other villages. Each cell shows the value of Fst x 1000. (D) Estimated effective migration surface plot within Sardinia based on 181 Sardinians with all four grandparents born in the same location. Abbreviations of subpopulations are: (Towns within Ogliastra) Arz, Arzana; Bar, Barisardo; Ilb, Ilbono; Gai, Gairo; Loc, Loceri; Lan, Lanusei; Tor, Tortoli; Vgs, Villagrande Strisaili, (Provinces) Cag, Cagliari; Car, Carbonia; Cmp, Campidano; Nuo, Nuoro; Olb, Obliatempio; Ori, Oristano; Sas, Sassari.

**Figure 3:**
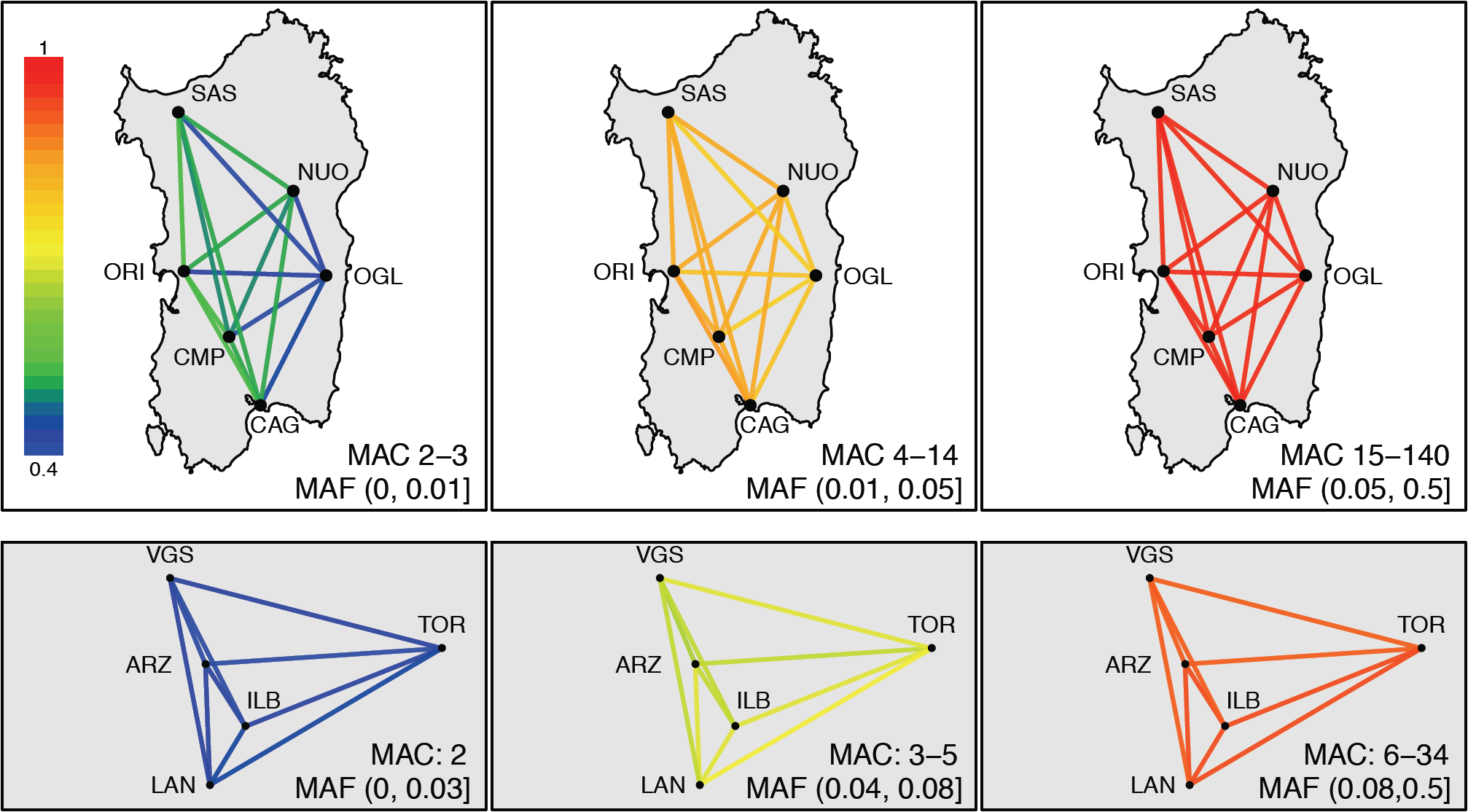
Allele-sharing across the island (top) and within Ogliastra (bottom) as a function of allele frequencies. Allele-Sharing between a pair of population is defined as the ratios of the probability that two randomly drawn carriers of the allele of a given minor allele frequency are from different populations, normalized by the panmictic expectation, and visualized here by the color of the lines connecting two populations. Island-wide analysis used only subpopulations with at least 70 individuals, and the minor allele counts of each of 1 million randomly selected variants were downsampled to 140 chromosomes. Within Ogliastra analysis used only subpopulations with at least 17 individuals, and the minor allele counts were downsampled to 34 chromosomes.

### Sardinia as an isolated Mediterranean population

To assess Sardinia in a broader context of the Mediterranean, we created a merged dataset of Sardinians and other previously analyzed Mediterranean and North African populations from the Human Origins Array (HOA) (Lazaridis et al. 2014). The Mediterranean Sea emerges as the most important feature affecting genetic structure in this data. PCA shows a one-dimensional isolation-by-distance configuration around the Mediterranean, from North Africa through the Near East and then towards Iberia (Novembre and Stephens 2008, Henn et al. 2012, Botigue et al. 2013, Paschou et al. 2014) (**Figure 4A,**). The effective migration surface shows the Mediterranean Sea as having low effective migration (**Figure 4B**), isolating Sardinia from neighboring mainland populations, with stronger isolation between Sardinia and North Africa than mainland Europe (**Figure 4B**).

**Figure 4:**
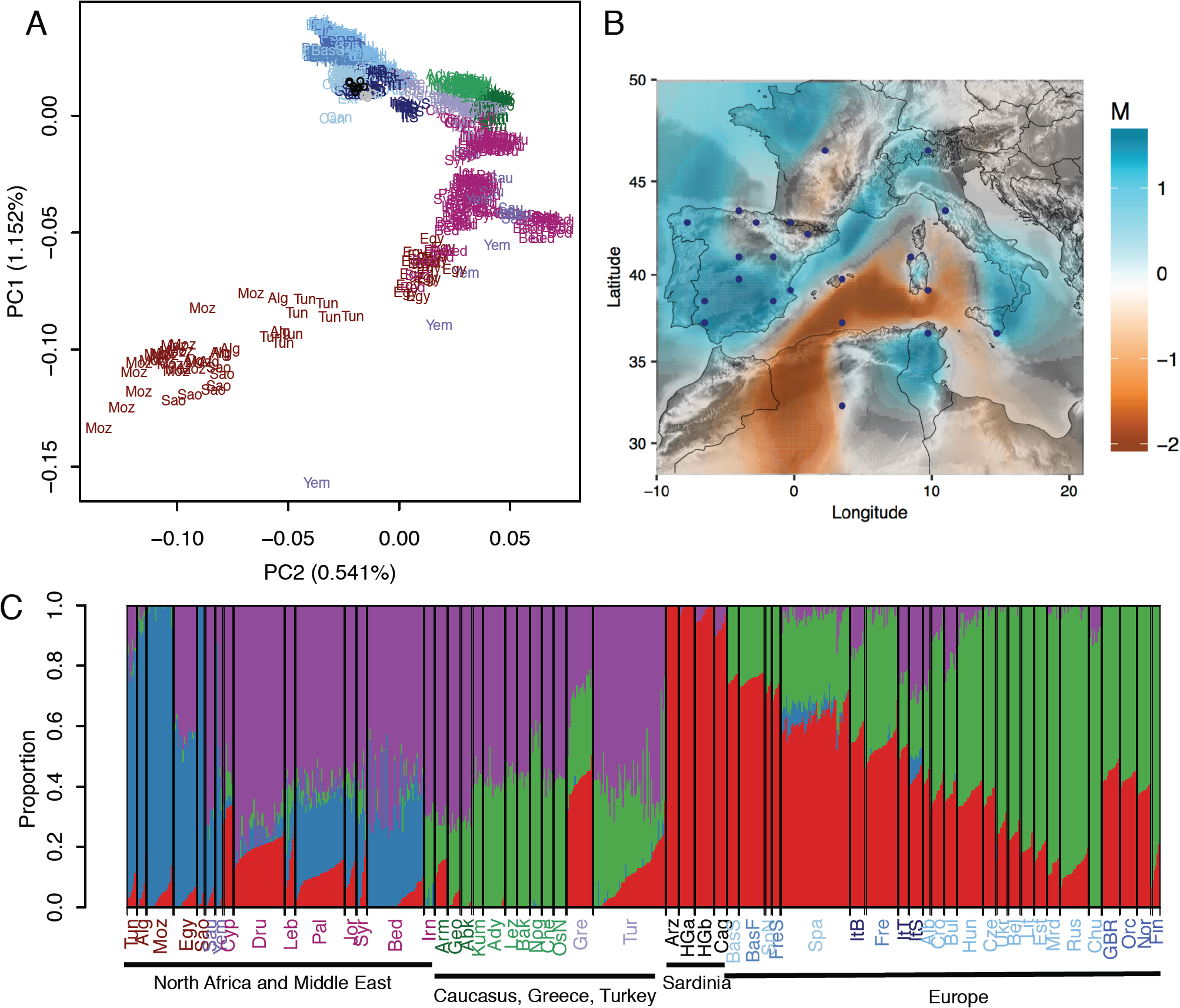
Population structure relative to mainland Europeans. (A) Top two components of PCA of the merged dataset of Sardinia and Human Origins Array data. Only populations from North Africa, Middle East, Caucasus, and Europe from the Human Origins Array data were included. Ten randomly selected Arzana and Cagliari individuals were used to represent the Gennargentu region and broader Sardinia, respectively. (B) Estimated effective migration surface result for the pan-Mediterranean analysis. (C) Admixture results at K = 4, which has the lowest cross-validation error in analysis from K = 2 to K = 15. See Figure S3 for the full result. The Sardinians consistently form a distinct cluster across Ks. Populations labels are color coded by major sub-continental regions. Abbreviations are: (North Africa) Sao, Saharawi; Moz, Mozabite; Alg, Algerian; Tun, Tunisian; Egy, Egyptian; (Middle East and Arabian Peninsula) Yem, Yemen; Sau, Saudi Arabian; Bed, Bedouin; Pal, Palestinian; Jor, Jordanian; Syr, Syrian; Dru, Druze; Leb, Lebanese; Irn, Iranian; Cyp, Cypriot; Tur, Turkey; (Caucasus) Arm, Armenian; Geo, Georgian; Abk, Abkhasian; Nog, Nogai; Ady, Adygei; OsN, North Ossetian; Kum, Kumyk; Che, Chechen; Bak, Balkar; Lez, Lezgin; (Europe) Gre, Greece; HGa, HGDP Sardinian; HGb, HGDP Sardinian; Cag, Cagliari; Arz, Arzana, ItS, Sicilians; ItyS, Italian South; ItT, Tuscan; ItB, Bergamo; Can, Canary Islands; Spa, Spanish; SpN, Spanish North; BasS, Spanish Basque; FreS, French South; Fre, French; BasF, French Basque; Alb, Albanian; Bui, Buglarian; Cro, Croatian; Hun, Hungarian; Chu, Chuvash; Cze, Czech Republican; Ukr, Ukranian; Bel, Belarusian; Mrd, Mordovian; Rus, Russian; Lit, Lithuanian; Est, Estonian; GBR, British Great Britain; Orc, Orcadian; Nor, Norwegian; Fin, Finnish.

As an alternative visualization of pan-Mediterranean population structure, an analysis using the ADMIXTURE software inferred four ancestral components, with one component associated primarily with Sardinians and Southern Europeans (“red”), and remaining components corresponding to North African (“blue”), Middle East and Caucasus (“purple”), and Northern Europeans (“green”) (**Figure 4C;** see **Figure S3** for results at other values of K). The Arzana individuals contained 100% of this red component and Sardinians from Cagliari contained 93% of this red component.

The isolation of Sardinia is especially evident in patterns of rare allele sharing. Using a doubleton sharing statistic, we also find that sharing between Sardinia and other mainland populations is small (normalized sharing ratio typically between 0.03 to 0.25), lower even than that between continental populations (e.g. approximately 0.3 to 0.7 between African and East Asian samples) (**Figure S4**). Within Sardinia, Arzana again shows evidence of being more isolated, with low sharing of alleles to the mainland (**Figure S4**). One interesting observation is a slightly elevated sharing between Sardinians and some non-European populations (LWK, PJL) and admixed Latino populations (MXL, PUR, CLM). This is particularly true for the non-Gennargentu Sardinian samples (**Figure S4**). The sharing with admixed Latino populations may reflect variants shared between Sardinian ancestors and ancestral European sources to admixed Latino populations (e.g. Iberian populations).

### Time-scale of divergence from mainland populations and population size history

While the relative isolation of Sardinia is apparent, the time-scale of the divergence is unclear and can be estimated using whole-genome sequence data. Here we used the coalescent-based MSMC method (Schiffels and Durbin 2014) and an analysis of the joint site frequency spectrum (SFS) using fastsimcoal (Excoffier et al. 2013) as two complementary approaches to estimate divergence times. We applied both methods with the 1000 Genomes’ Utah (CEU) and Tuscan (TSI) populations as reference outgroups. Using MSMC, we found the cross-coalescent rates between Sardinia and TSI or CEU decline to 0.5 approximately 327 generations ago (0.25-0.75 range = 280 to 539 gen for Sardinia-CEU, 280 to 428 for Sardinia-TSI) which is earlier than divergence between TSI and CEU at approximately 190 generations ago (0.25-0.75 range = 110 to 276 gen) (**Figure 5B**). The analysis of the joint site frequency spectra corroborates this result, with Sardinians diverging from Northern Europeans 349 generations ago (95% C.I. = 89 to 403 gen), while TSI diverging from Northern Europeans 202 generations ago (95% C.I. = 55 to 374 gen) (**Table S3, Figure S5**). These methods also support that Sardinia has had low long-term effective population sizes, and lacks the signature of strong population growth typical of mainland European populations (**Figure 5A**, **Table S3**).

**Figure 5:**
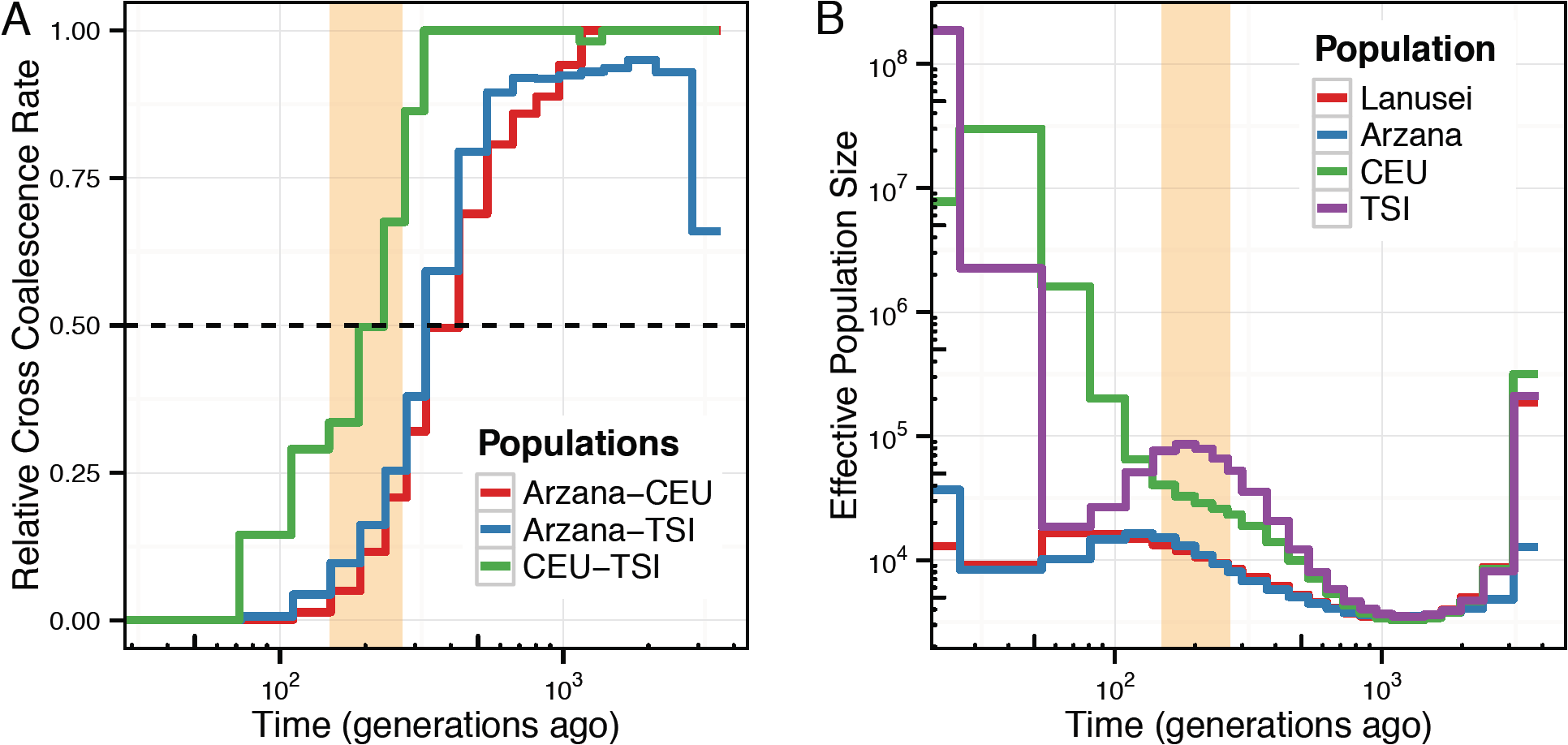
Coalescent-based inference of demographic history using MSMC. (A) Inferred relative cross coalescence rate between pairs of populations through time, based on 4 haplotypes each from Arzana, CEU, and TSI. (B) Population size history inference based on 8 haplotypes of high-coverage individuals from each of Lanusei (LAN), Arzana (ARZ), 1000 Genomes CEU and tSi. The shaded box denotes roughly the Neolithic period, approximately 4,500 to 8,000 years ago, converted to units of generations assuming 30 years a generation.

### Sardinia in relation to other Mediterranean populations, especially the Basque

Due to its smaller long-term effective population size (**Figure 5A**), Sardinia is expected to have undergone accelerated rates of genetic drift. To correct for this when measuring similarity to other mainland populations, we used “shared drift” outgroup-f3 statistics (Raghavan et al. 2014), which are robust to population-specific drift. Using this metric, we find the Basque are the most similar to Sardinia, even more so than neighboring mainland Italian populations such as Tuscany and Bergamo (**Figure S6A, S6B**). This relationship is corroborated by identity-by-descent (“IBD”) tract length sharing, where among mainland European populations, French Basque showed the highest median length of shared segments (1.525 cM) with Arzana (**Figure S7**). We also tested the affinity between Sardinians and Basque with the D-statistics of the form D(Outgroup, Sardinia; Bergamo or Tuscan, Basque). In this formulation, significant allele sharing between Sardinia and Basque, relative to sharing between Sardinia and Italian populations, will result in positive values for the D-statistic. We find that Sardinia consistently showed increased sharing with the Basque populations compared to mainland Italians (|Z|> 4; **Figure S6C**), and the result was stronger when using the Arzana than Cagliari sample (D_ARZ_ = 0.008 and 0.0096, D_CAG_ = 0.0072 and 0.0087 for French Basque and Spanish Basque, respectively). In contrast, sharing with other Spanish samples in our dataset was generally weaker and not significant (|Z| < 3.5; **Figure S6C**), suggesting the shared drift with the Basque is not mediated through Spanish ancestry.

The ADMIXTURE and PCA analyses above (**Figure 4**) suggest that Sardinian samples, such as those outside of Ogliastra, may show some evidence of admixture with mainland sources, consistent with previous reports (Moorjani et al. 2011, Loh et al. 2013, Hellenthal et al. 2014). For example, Cagliari individuals demonstrated ~7% of a non-Sardinian (“purple”) component that is found in substantial fraction among extant individuals from Southern Europe, Middle East, Caucasus, and North Africa. To assess this in detail we used the f3-test for admixture (Patterson et al. 2012) and found none of the Sardinian populations showed any evidence of admixture. In contrast, mainland Europeans, particularly Southern Europeans, show evidence of admixture from Near East and sub-Saharan Africa (**Figure S8**). Because f3-based tests for admixture may lose power when applied to populations that have experienced extensive drift post-admixture (Patterson et al. 2012), we also used a complementary LD-based approach (ALDER, Loh et al. 2013) to test each Sardinian population for admixture. Using this approach, a number of Sardinian populations, particularly those outside of Ogliastra, are inferred to be admixed (**Table 1, Table S4**). The inferred source populations are typically a mainland Eurasian population and a sub-Saharan African population. The admixture proportions range from 0.9% to 4% of sub-Saharan ancestry by the f4 ratio test (Patterson et al. 2012) (**Table 1, Table S4**) with estimated admixture dates of approximately 59-109 generations.

**Table 1:**
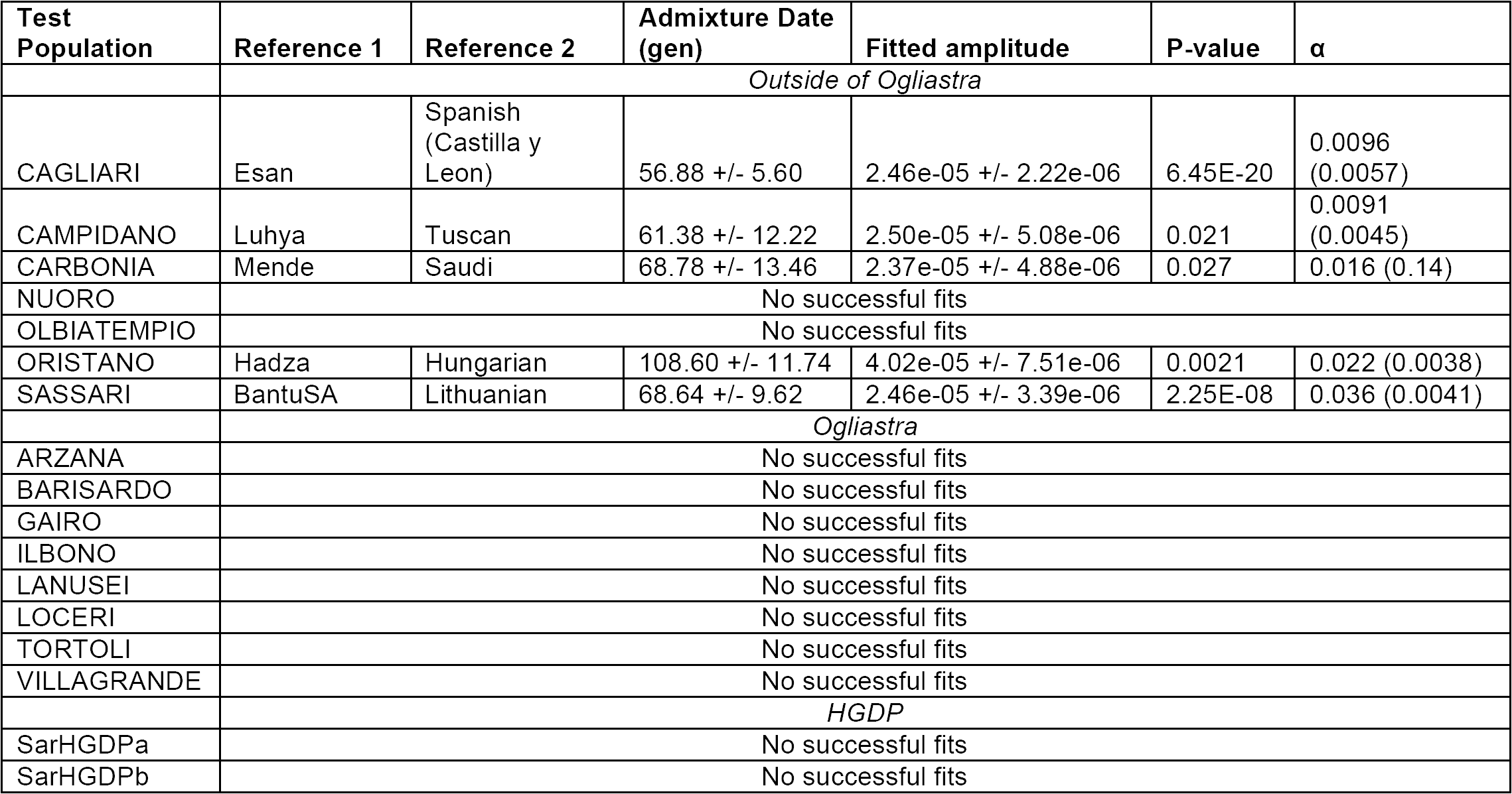
Evidence of admixture as inferred by ALDER. P-value has been corrected for multiple hypothesis testing (both number of pairs of source populations and analysis-wide number of test populations). α is the estimated admixture proportion of the sub-Saharan ancestry by the f4 ratio test, using Finnish and Chimp as the outgroups (except for Carbonia, where Esan and Chimp were used and the result was consistent with 0% admixture).

### Elevated Neolithic and pre-Neolithic ancestry in Gennargentu region

Ancient DNA studies have shown that across the autosome, Sardinians exhibit higher levels of Neolithic Farmer ancestry compared to other mainland Europeans (Lazaridis et al. 2014, Haak et al. 2015). However, because previous representative samples from Sardinia were from the HGDP, which is limited in sample size, we sought to revisit the questions using using our dataset and addressing within-island variation.

We generally find that Sardinians have the highest observed levels of shared drift with early Neolithic farming cultures and low levels of shared drift with earlier hunter-gather samples (**Figure 6A, Figure S9, Table S5**), consistent with previous reports. Surprisingly though, when examining allele sharing within Sardinia, we found that both ancient Neolithic farmer ancestry and pre-Neolithic ancestry are enriched in the Gennargentu-region. First, we find that shared drift with Neolithic farmers and with pre-Neolithic hunter-gatherers is significantly correlated with the proportion of “Gennargentu-region” ancestral component estimated from ADMIXTURE analysis, while shared drift with Steppe pastoralists has a weak negative correlation with Gennargentu-region ancestry (**Figure 6B**). Second, using supervised estimation of ancestry proportion based on aDNA (Haak et al. 2015), we estimate higher levels of Neolithic and pre-Neolithic ancestries in the Gennargentu region and higher levels of Steppe Pastoralist ancestry outside the region (**Figure S10**). Finally, calculations with Patterson’s D-statistics of the form D(Outgroup, Ancient, Ogliastra, Non-Ogliastra) also support increased sharing with Neolithic and pre-Neolithic individuals, but not post-Neolithic individuals from the Steppe, in the Ogliastra samples (D = -0.0037 and -0.0042, |Z| = 7.4 and 7.9 when aDNA sample = Stuttgart and Loschbour, respectively; D = -0.0009, |Z| is not significant, when aDNA sample = Yamnaya).

**Figure 6:**
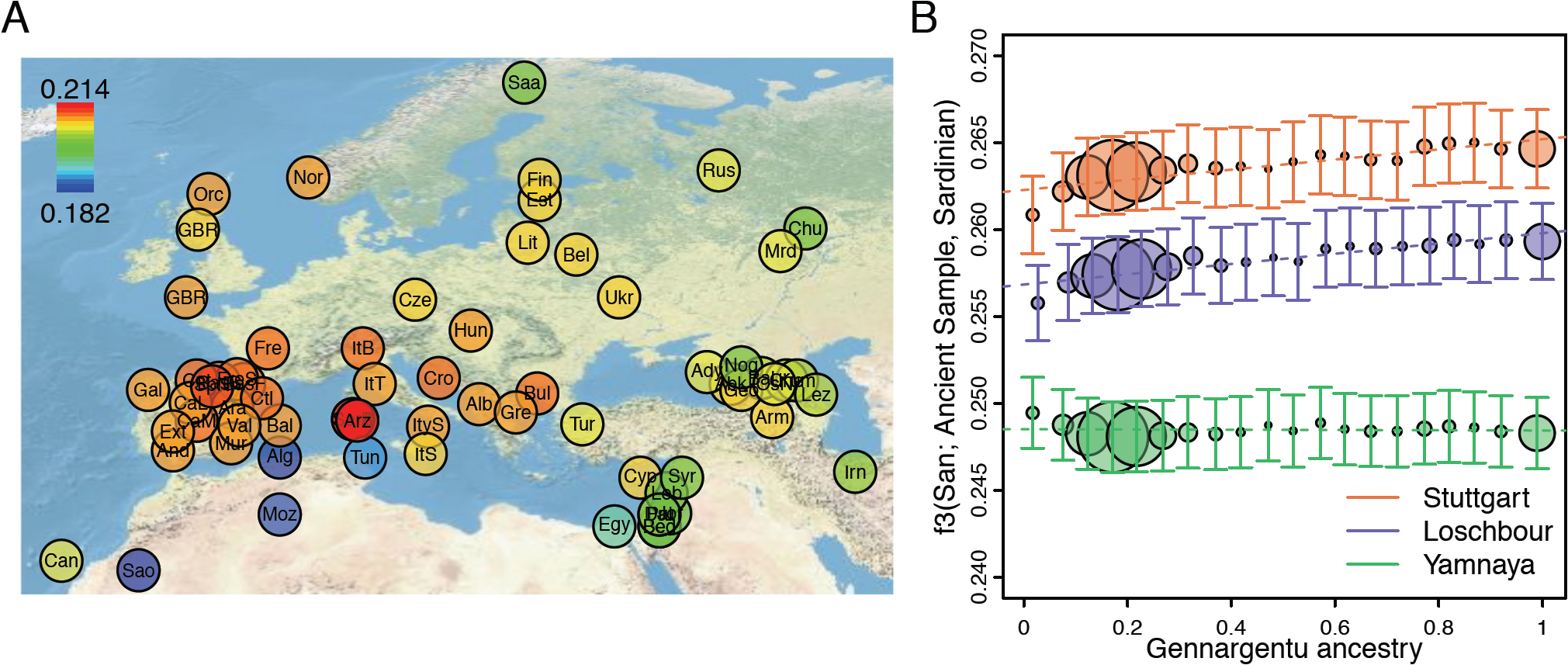
Similarity of ancient samples to populations across Europe and within Sardinia. (A) Outgroup f3 statistics of the form f3(San; Stuttgart, X), where X is a population across the merged dataset of Sardinia and Human Origins Array data. Higher f3 values suggest larger shared drift between a pair of populations. Population abbreviations are the same as Figure 4. (B) Outgroup f3 statistics of the form f3(San; Ancient, Sardinian) across Sardinian samples binned in steps of 5% of “Arzana” ancestry estimated in Figure 2B. The increase of outgroup f3 statistics as function of ancestry is positive for Stuttgart and Loschbour (0.00292 and 0.00295, respectively), and slightly negative for Yamnaya (-7.4e-5). Ancient samples used include a reference Neolithic farmer individual (Stuttgart, orange), a reference pre-Neolithic hunter-gather individual, (Loschbour, blue), and a reference Steppe population(Yamnaya, green) from the merged dataset with Haak et al. (Methods). Sizes of the circle are proportional to the number of samples per bin (max N = 281 per bin).

Together, these results suggest that while at the regional-level Sardinia appears to harbor the highest amounts of Neolithic farmer ancestry and very little of the pre-Neolithic hunter-gatherer or Bronze Age pastoralists ancestries, there exists within-island variation. Specifically, we find that with increasing level of isolation (represented by increasing level of the Gennargentu ancestry), there are greater Neolithic farmer and pre-Neolithic hunter-gatherer ancestry, while the less isolated Sardinians have a stronger signal of ancestry from the Steppe pastoralist source.

### Sex-biased demography in prehistoric Sardinia

As described in the Introduction, the discrepancy between Y chromosome data and the autosomal signature of Neolithic ancestry in Sardinia raises the questions of whether sex-specific processes have occurred in the history of Sardinia. The availability of genome-wide data allows for a novel evaluation of sex-specific processes by using X versus autosome comparisons (Hammer et al. 2008, Bustamante and Ramachandran 2009, Keinan et al. 2009, Goldberg et al. 2016).

To carry out such comparisons, we first use ADMIXTURE and contrast the inferred Gennargentu-region ancestry on the X chromosome versus the autosome. Intriguingly, on average, we find a higher proportion of the Gennargentu-region ancestry (“red” component in **Figure S11**) on the X-chromosome (37%) than on the autosome (30%, *P* < 1e-6 by permutation). As the Gennargentu-region ancestry is correlated with more ancient, Neolithic or pre-Neolithic ancestries rather than Bronze Age ancestries (**Figure 6B**), this finding suggests sex-biased processes in which more females than males carried the non-Steppe ancestries.

To more directly delineate this result using aDNA samples, we contrasted D statistics computed separately on the X chromosome and on the autosome using several aDNA samples (Neolithic farmer individuals such as Stuttgart and NE1 (Gamba et al. 2014, Lazaridis et al. 2014), and pre-Neolithic hunter-gatherer individuals such as Bichon and Loschbour (Lazaridis et al. 2014, Jones et al. 2015)). Specifically, we computed D(Outgroup, Ancient, Sardinians, CEU), which would lead to a negative value if there is increased allelic sharing between Sardinians and the ancient individuals, relative to mainland Europeans (represented by CEU). The presence of sex-biased demography would cause D statistics on the X chromosome to be significantly different from those calculated on the autosomes.

Using this approach, we find an excess of Neolithic, and to a lesser extent, pre-Neolithic ancestries in Sardinians on the X chromosome relative to the autosomes (**Figure 7,** D_x_ < D_A_; *P* < 0.0001 by bootstrapping). In contrast, we recapitulate the expected pattern of excess Neolithic ancestry and depletion of pre-Neolithic ancestry in Sardinia relative to mainland Europeans on the autosome. The excess of ancestries represented by Neolithic and pre-Neolithic individuals on the X versus autosomes is robust to using different outgroups (**Figure S12**), comparing the X chromosome only to the similarly gene-dense chromosome 7 (**Figure S13**), and testing with different ascertainment panels (**Figure S14**). Using other 1000 Genomes populations in place of the Sardinians as controls showed little difference between X chromosome and the autosome (**Figure 7**). These results demonstrate that sex-biased process occurred in the founding of Sardinia, which is above and beyond any putative sex-biased processes on the mainland (Goldberg et al. 2016).

**Figure 7:**
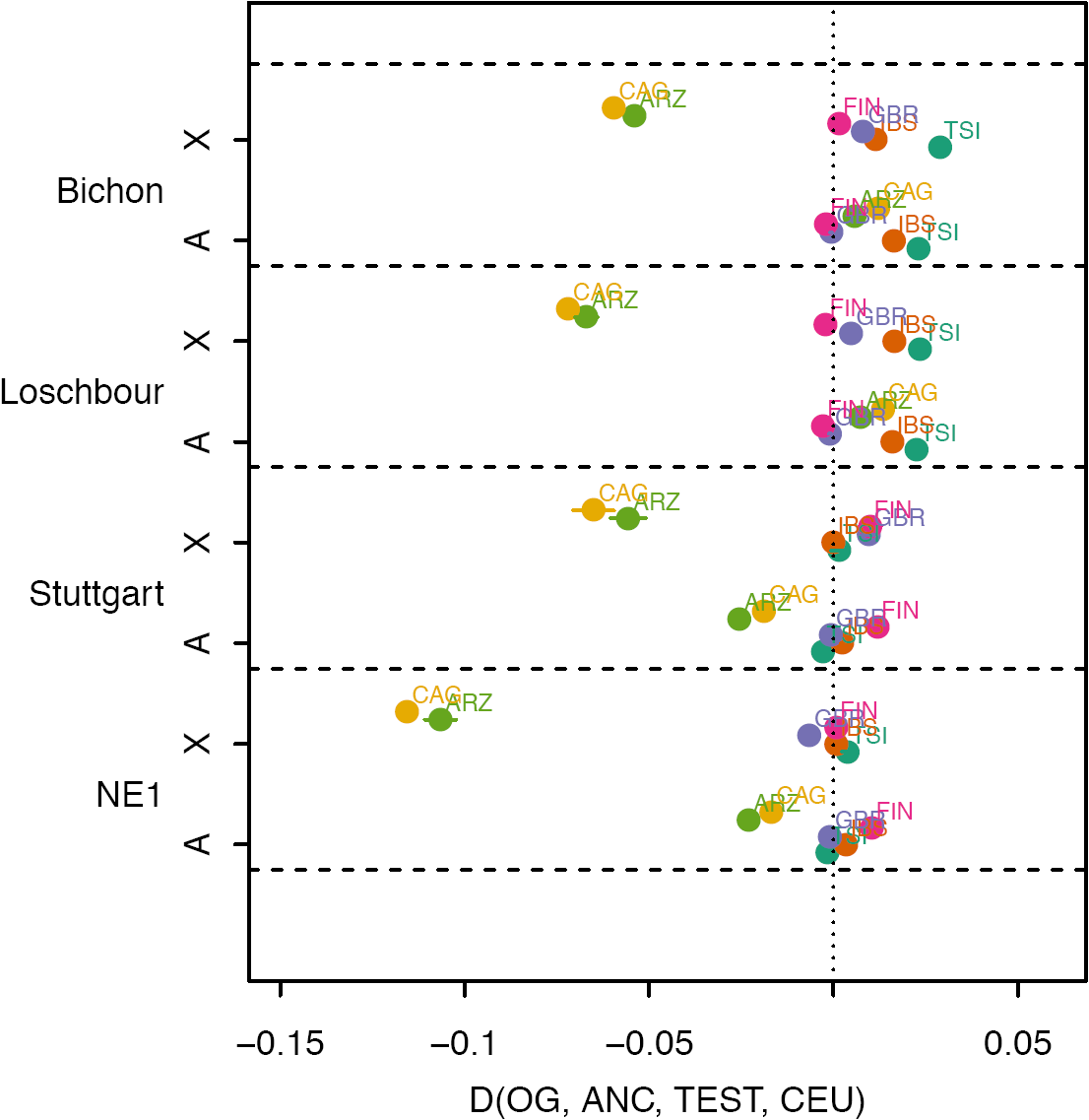
Evidence of sex-biased demography by contrasting chr X vs. autosome. D statistics of the form D(Outgroup, Ancient, Test, CEU) were computed separated using X chromosome and autosomal data (Method). Ancient samples tested were Neolithic farmer individuals NE1 and Stuttgart, and pre-Neolithic hunter-gatherer samples Loschbour and Bichon. Test populations include each of the Sardinian subpopulations (Cagliari and Arzana are visualized here), as well as TSI, IBS, and FIN from 1000 Genomes. Outgroup is YRI from 1000 Genomes. If there is no sex-biased demography relative to CEU, the D statistics contrasting X and autosome should be approximately equal, as observed generally for the 1000 Genomes populations.

## DISCUSSION

We investigated the fine-scale population structure and demography of the people of Sardinia using a whole-genome sequenced dataset of 3,514 Sardinians. The genotype calling procedure for this dataset leveraged extensive sharing of haplotypes across distantly related Sardinians to produce a high quality call set (Sidore et al. 2015), and we cross-referenced the genetic dataset with detailed self-reported ancestry and geographical labeling that goes back two generations. By focusing on a subset of 1,577 unrelated Sardinians and intersecting them with the 1000 Genomes, HGDP, and Human Origin Array reference sets we were able to analyze the population genetics of Sardinia populations in greater detail. We were able to confirm a number of major features of previous analyses: the differentiation of Sardinia from mainland populations, the presence of high Neolithic farmer ancestry in Sardinia, and the presence of a small amount of sub-Saharan African admixture. Further, our analyses provide more detail regarding the isolation between Sardinia and the mainland. Our analysis of cross-coalescent rates suggest the population lineage ancestral to modern-day Sardinia was effectively isolated from the mainland European populations approximately 330 generations ago. This estimate should be treated with caution, but corresponds to approximately 9,900 years ago assuming a generation time of 30 years and mutation rate of 1.25x10^−8^ per basepair per generation.

We documented fine-scale variation in the ancient population ancestry proportions across the island. The most remote and inaccessible areas of Sardinia–the Gennargentu Massif covering the central and eastern regions, including the present day province of Ogliastra, would have been the least exposed to contact with outside populations (Morelli et al. 2000, Pistis et al. 2009, Ghirotto et al. 2010). That pre-Neolithic hunter-gatherer and Neolithic farmer ancestry are enriched in this region of isolation suggests that the early populations of Sardinia were an admixture of the two ancestries, rather than the pre-Neolithic ancestry arriving via later migrations from the mainland. However, it remains to be seen whether this admixture principally occurred on the island or on the mainland prior to a Neolithic era influx to the island.

We also found Sardinians show an impressive signal of shared ancestry with the Basque, in terms of identity-by-descent tracts and the outgroup f3 shared-drift metric. Such a connection is consistent with long-held arguments of a connection between the two populations, including claims of Basque-like non-Indo-European language words among Sardinian placenames (Blasco Ferrer 2010). More recently the Basque have been shown to be enriched for Neolithic farmer ancestry (Lazaridis et al. 2014, Gunther et al. 2015) and Indo-European languages have been associated with Steppe population expansions in the Bronze Age (Allentoft et al. 2015, Haak et al. 2015). These results support a model in which Sardinians and the Basque may both retain a legacy of pre-Indo-European, Neolithic ancestry (Gunther et al. 2015).

We also examined possible sources of African admixture to Sardinia. Prior to our studies, there have been reports based on the HGDP Sardinians of a minor proportion (0.6% to 2.9%) of sub-Saharan admixture (Moorjani et al. 2011, Loh et al. 2013) and a multi-way admixture involving an African source (Hellenthal et al. 2014). In light of the close geographical proximity of Sardinia and North Africa, as well as the substantial admixture proportion from North Africa in Southern Europe (Botigue et al. 2013), we tested for admixture using modern North African reference populations included in the Human Origins Array data (Tunisia, Algeria, Mozabite, Egypt, and Saharawi), and found the best proxy for African admixture is sub-Saharan African populations, rather than Mediterranean North African populations, and we inferred the date of admixture as approximately 1,800-3,000 years ago (assuming 30 years per generation). The lack of a strong signal of North African autosomal admixture may be due to inadequate coverage of modern North African diversity in our reference sample. Alternatively, it may be due to a poor representation of ancestral North Africans. Present-day North African ancestry reflects large-scale recent gene flow during the Arab expansion (~1,400 years ago (Henn et al. 2012)). The sub-Saharan African admixture observed in the non-Ogliastra samples could be mediated through an early influx of migrants from North Africa prior to the Arab expansion, for example during the eras of trade relations and occupations from the Phoenicians, Carthaginians, and Romans (~700 B.C.- ~200 B.C.; (Dyson and Rowland 2007)).

The high frequency of particular Y-chromosome haplogroups (particularly I2a1a2 and R1b1a2) that are not commonly affiliated with Neolithic ancestry is one challenge to a model in which Sardinian principally has Neolithic ancestry. Whether such haplogroup frequencies are due to simple genetic drift and/or a signal of sex-biased demographic processes has been an open question. By carrying out X versus autosome comparisons we uncovered evidence of sex-biased patterns of ancestry in Sardinia, and found an enrichment of Neolithic and pre-Neolithic ancestry on the X-chromosome across all groups on the island. Using X/autosome comparisons to study sex-biased demography has been challenging, due to many distinct factors that can affect the patterns of variation (Hammer et al. 2008, Bustamante and Ramachandran 2009, Keinan et al. 2009). Here, we note that we tested whether the enrichment of Neolithic ancestry on the X is above and beyond what is observed in CEU, and this provides an internal control against hidden confounders. We also replicated our results by comparing chromosome X specifically to only chromosome 7, which is the most closely matched autosome in terms of number of basepairs sequenced and the known gene density in the human reference genome (Scherer 2010).

A remaining challenge is synthesizing these results to provide a comprehensive model of Sardinian population history, including mechanisms for the sex-biased ancestry we observe. While this involves uncertainty, one model for the complex patterns of genetic variation we observe is that the present population is the result of an early pre-Neolithic contribution (marked by the I2a1 lineage and the enrichment of HG ancestry in the isolated Gennargentu region) and a larger contribution of Neolithic farmer ancestry, most likely accentuated by the obsidian trade between mainland Europe and Sardinia during the early Neolithic. The data are also consistent with a subsequent male-biased influx of Steppe ancestry after the Bronze Age, especially outside of the Gennargentu region. Such gene flow would have diluted the autosomal Neolithic and pre-Neolithic components relative to the X chromosome, resulting in the pattern of sex-biased allele sharing we observe. This model predicts that the effect of sex-biased allele sharing would be larger for non-Gennargentu populations than Gennargentu populations, since the impact of Steppe ancestry is stronger outside of Ogliastra (**Figure S10**). Our results from the D-statistic analysis is consistent with this prediction (D**_CAg,x_**-D**_CAg,a_** = -0.046 to -0.139, D_ARZ,X_-D_ARZ,A_ = -0.030 to -0.084, *P* = 0.0012 to 0.0015). Also consistent with this model is the higher frequency of R1b1a2 outside of Ogliastra (19%) than in Ogliastra (9.4%) (Zei et al. 2003, Francalacci et al. 2013, Haak et al. 2015). Note that while R1b1a2 was previously interpreted as a marker for pre-Neolithic ancestry, current aDNA data suggest it may be a better proxy for the Bronze Age expansion (Haak et al. 2015). This model is also consistent with recently described evidence of male-biased migration from the Steppe among mainland Europeans at large (Goldberg et al. 2016).

Such a model, however, does not immediately explain the high prevalence of the I2a1a1 haplogroup in Sardinia given the predominantly Neolithic farmer ancestry of the population. As reviewed above, the I2a1a1 haplogroup appears to be pre-Neolithic and is at high frequency in Sardinia (~40%). The frequency of the I2a1a1 haplogroup could have increased within a general Neolithic farmer background either during bottlenecks associated with early migrations to Sardinia or by drift on the island. However, neutral forward simulations based on the population size history estimated here (**Figure 5A**) showed that I2a1a1 haplogroup is unlikely to drift to such high frequency (**Figure S15**). Therefore, additional sex-biased processes, such as male-biased drift mediated by small effective sizes for males may have been acting as well. Small effective sizes for males can arise due to polygyny, patrilineal descent rules, or transmission of reproductive success (Heyer et al. 2012).

Further investigation is ultimately necessary to establish whether this model with male-mediated Steppe influx is correct, or whether alternative models need further consideration. For example, processes that would lead to female-biased migration from the Neolithic sources could be consistent with the enrichment of Neolithic ancestry on the X and elevated pre-Neolithic ancestry on the Y; or one might consider a model in which a yet unsampled sub-population during the Neolithic carried high proportions of the I2a1a haplogroup. Our results here do make clear that the peopling of Europe is more complicated than portrayed in basic 3-component ancestry model and more attention needs to be paid to understanding sex-biased processes in the history of European populations. Future studies focusing on inferring sex-stratified demographic parameters on the basis of the demographic history presented here and ancient DNA data from Sardinia and Southern Europe will be illuminating, especially as systems of mating and dispersal may have shifted alongside modes of subsistence (Wilkins and Marlowe 2006).

For the purposes of understanding the evolution of complex quantitative trait evolution in Sardinian history, the results suggest that while Sardinia has clearly had influence from pre-Neolithic sources and contact with Steppe populations, the demographic history is one of substantial isolation and abundant Neolithic ancestry relative to the mainland. For traits with a strong component of sex-linked QTL our results encourage accounting for the sex-biased processes detected here. The relatively constant size of Sardinian populations predicts there will be less of an influx of rare variants (Keinan and Clark 2012), as well as an increase of homozygosity, relative to other expanding populations. These two factors may increase the impact of dominance components of variation (Joshi et al. 2015), and reduce the allelic heterogeneity of complex traits (Lohmueller 2014, Simons et al. 2014, Uricchio et al. 2016). Armed with a better understanding of Sardinian prehistory and demographic events, we anticipate a more nuanced understanding of complex trait variation and disease incidences in Sardinia. The affinity to Neolithic farmer populations (and to a lesser extent, the pre-Neolithic hunter-gatherer populations) also means Sardinia is a potential reservoir for variants in these ancient populations that may have been lost in mainland Europeans today. Thus a phenome-wide association study to these farmer- or hunter-gatherer-specific alleles may also shed light on the evolution of traits from the past to the present.

## ACKNOWLEDGEMENTS

This study was funded in part by the National Institutes of Health, including support by National Human Genome Research Institute grants HG005581, HG005552, HG006513, HG007089, HG007022, and HG007089; by National Heart Lung and Blood Institute grant HL117626; by the Intramural Research Program of the NIH, National Institute on Aging, with contracts N01-AG-1-2109 and HHSN271201100005C; National Institute of General Medical Sciences with GM108805 and T32NS048004; National Institute of Neurological Disorders and Strokes with T32NS048004; and an NSRA postdoctoral fellowship F32GM106656. This research was also supported by Sardinian Autonomous Region (L.R. no. 7/2009) grant cRP3-154; by PB05 InterOmics MIUR Flagship Project; by grant FaReBio2011 “Farmaci e Reti Biotecnologiche di Qualita”.). H.A. was supported by an NSF Graduate Research Fellowship. We also would like to thank Iosif Lazaridis, Nick Patterson, Sohini Ramanchandran and Robert Brown for discussion and technical assistance, as well as members of the Novembre and Lohmueller lab for constructive comments regarding this research.

## AUTHOR CONTRIBUTIONS

F.C. G.R.A., D.S., and J.N. conceived of the study; C.W.K.C., C.S., D.S., F.C. G.R.A., J.N. designed the study; C.W.K.C., J.H.M., C.S., H.A. performed the analyses; C.W.K.C., J.H.M., C.S., K.E.L., G.R.A., D.S., F.C., J.N. interpreted the data; C.S., M.Z., M.P., F.B., A.M., G.P., M.S., A.A., G.R.A., D.S., F.C. contributed in the data collection and initial preparation for population genetic analysis. C.W.K.C., J.H.M., C.S., H.A., D.S., F.C., and J.N. wrote the paper with input from all coauthors.

## METHODS

### Cohort description

We included in this study individuals from the SardiNIA/Progenia longitudinal study of aging (Pilia et al. 2006, Sidore et al. 2015) based in the Ogliastra region and from the case-control (CSCT) studies of Multiple Sclerosis (Sanna et al. 2010) and Type 1 Diabetes (Zoledziewska et al. 2009) based on a sampling effort across the general population of Sardinia. For the case-control study cohort, we required individuals to have at least three Sardinian grandparents. All participants gave informed consent, with protocols approved by institutional review boards for the University of Cagliari, the National Institute on Aging, and the University of Michigan.

### Whole-genome sequenced Sardinian dataset

The dataset includes 3,514 individuals sequenced at low-coverage (average coverage 4.2x) and 131 high-coverage individuals (average coverage 36.7x). 2,090 individuals belong to the SardiNIA cohort, the remaining 1,424 are derived from the case-control study. A subset of 2,120 low-coverage individuals were previously described (Sidore et al. 2015). The additional 1,394 individuals and the 131 high coverage individuals were aligned, recalibrated and quality checked using the same criteria(Sidore et al. 2015) to guarantee sample uniformity. Variant calling was performed using GotCloud as described in (Sidore et al. 2015), which also include a step of genotyping refinement with Beagle (Browning and Browning 2009) to increase genotype accuracy in the low-coverage individuals.

To generate reliable calls for the X chromosome we performed standard variant calling to generate genotype likelihoods for each individual genotype. After that, we modified the likelihoods of male individuals to reduce the likelihoods of heterozygous genotypes (heterozygotes GL set to 500) and ran genotype refinement as described for autosomal markers. We then converted the male diploid genotypes generated by Beagle into haploid genotypes.

To process the high-coverage data, we created a pileup of raw sequence reads using samtools (“samtools mpileup”), filtering out bases with a base quality score < 20 and reads with a map quality score < 20. We then used the bcftools variant caller (“bcftools call”) in conjunction with a custom script from the msmc github repository (https://github.com/stschiff/msmc-tools) to call single nucleotide polymorphisms. We then phased each individual with SHAPEIT2, using the 1000 Genomes Phase 3 data reference panel as provided on the SHAPEIT2 website (https://mathgen.stats.ox.ac.uk/impute/impute_v2.html#reference).

### Filtering individual samples by poor sequencing quality and relatedness

We aimed to identify a set of unrelated and reliably genotyped individuals for most of the population genetic analysis presented in this paper. Therefore we first excluded individuals with low-quality genotypes on the basis of an excess amount of genotypes with low confidence from the genotype refinement step. Specifically, examining the proportion of imputed genotypes with highest posterior genotype probability less than 0.9 (**Figure S16**), we found 8 outlier individuals with an excess amount (> 0.992) of low quality genotype calls. Manual inspections suggest the poor imputation quality is due to low coverage of these samples, and thus these samples were removed from down-stream analysis.

To prune the dataset for related individuals, we first extracted a subset of 153.7K SNPs with maximum pairwise *r*^*2*^ of 0.2 (pruned from a subset of 1.21M SNPs overlapping between the Sardinian whole genome sequence data and HapMap 3), we used PLINK to compute the genome-wide proportion of pairwise identity of descent sharing (pihat). The distribution of pihat (**Figure S17**) showed distinct modes corresponding to different degrees of relatedness, as well as extensive low level sharing (pihat < 0.1) between individuals, consistent with long-term isolation and endogamy. We opted to use a pihat cut-off of 0.07, and for each pair of individuals with pihat > 0.07, we preferentially removed the offspring if in a trio, or otherwise the individual appearing to be more related to the rest of the sample (by the sum of pihat in all other relationships with pihat > 0.07), until there were no relationships with pihat > 0.07 left. In total, we are left with 1,577 unrelated individuals, including 615 individuals from the SardiNIA sample and 964 individuals from the case-control cohort. Because the sampling design favored sampling of individuals from families, especially in the SardiNIA cohort, a large reduction in the sample size due to the relatedness filters is expected.

### Defining sample origin based on self-reported grandparental ancestry

Study participants were asked to self-report the location of residence of each of their parents and grandparents. We aimed to assign a 4-part ancestral origin to each sample based on these self-reports. We first categorize each geographical location by three levels of resolutions: (1) macro-regions (e.g. Sardinia, South Italy, France, Tunisia), (2) provinces within Sardinia (e.g. Cagliari, Sassari, Ogliastra), (3) town level (e.g. Arzana, Lanusei, Tortoli). For each parental lineage, if origin information is available for both grandparents and the parent, we preferentially use grandparental origin. If grandparental information is missing but parental information is available, we use the parental information but use the approximation that it represents the ancestry from both grandparents. If origin information is available for only one grandparent, we use that information and then assign the other grandparent origin as missing. Given the more detailed information provided by the participants of the SardiNIA project, we defined SardiNIA samples down to town resolution, but only define the case-control samples down to the province resolution unless noted otherwise.

After assigning a 4-part origin to each sample, we initially examined whether the distribution of genetic ancestry may differ based on these assignments. Empirically, we see that the genetic ancestry did not significantally differ for individuals having 4- or 3- parts of their ancestry coming from a particular location (**Figure S18**). It also did not differ significantly by whether the 4-part origin came from self-reported grandparental origin or self-reported parental origin (data not shown). However, we did observe in Admixture and PCA analyses that individuals with 2-part origin are more heterogeneous (**Figure S18** and data not shown). Thus for the majority of the downstream analysis we grouped individuals by a simple majority rule, requiring that individuals have at least 3 out of the 4-part origin from the same geographical location; otherwise the individual is called “split” and dropped from most analyses where discrete geographical labeling is used.

### Merging with other reference array datasets

For analyses investigating the genetic relationship between Sardinia and neighboring populations from mainland Europe, Middle East, and Northern Africa, we merged the Sardinia sequenced data with the Human Origin Dataset (Lazaridis et al. 2014). Out of the 600,841 SNPs released with the Human Origin Dataset, 489,747 were found in the Sardinian whole genome sequencing dataset. We then removed SNPs showing inconsistent alleles between the two datasets (485), as well as SNPs with greater than 2% missing genotypes in either dataset (14,525), and merged the two dataset over the remaining SNPs (474,737). The Human Origin Dataset contained no A/T or C/G SNPs, thus strand inconsistencies due to these transversion SNPs are not an issue. Finally, to avoid analyzing only variants polymorphic in Sardinia that could bias downstream f3 and D statistics, we assumed that all Sardinian individuals are homozygous for the reference allele at autosomal variants found in the Human Origin Dataset but not in the Sardinian genome sequencing dataset (104,664). Thus in total we have a merged dataset of 579,401 variants. Some basic information and summary statistics of the reference panels and populations used in the merge can be found in **Table S6**. For estimating mixture proportions in a 3-way model of European admixture, we merged our dataset with the version of Human Origin Array data published by ref. (Haak et al. 2015), which contains more ancient samples but fewer SNPs. We applied the same merging procedures as described above, and used a total of 345,822 SNPs for the analysis. Unless denoted specifically, the Human Origins merge is with the Lazaridis et al data.

### PCA and Fst

Sardinia-specific PCA was conducted using all unrelated individuals genotyped at SNPs found in HapMap 3 (International HapMap et al. 2010). For regional PCA, since a significant imbalance of sample sizes across populations may distort the PCA, a random subset of 10 unrelated Sardinians from Arzana and Cagliari were chosen to represent Sardinia and merged with the Human Origin dataset.

PCA analysis was performed using EIGENSTRAT version 5.0 after removing one SNP of each pair of SNPs with r^2^ ≥0.8 (in windows of 50 SNPs and steps of 5 SNPs) as well as SNPs in regions known to exhibit extended long-range LD (Price et al. 2008). In total 519K and 328K SNPs were used in the Sardinia-specific PCA and regional PCA, respectively.

Weir and Cockerham’s unbiased estimator of Fst was also calculated in pairwise fashion between populations grouped at the province level in Sardinia or grouped at the village level within Ogliastra using the same set of pruned SNPs.

### Admixture

Similar to the PCA analyses, Sardinia-specific Admixture analysis was conducted using all unrelated individuals genotyped at SNPs found in HapMap 3, while the regional analysis was conducted with sub-sampling of 10 Sardinians each with self-reported Cagliari and Arzana ancestries. Analysis was performed using Admixture version 1.22, following the recommended practice in the manual for LD filtering (removing one SNP of each pair of SNPs with r^2^ ≥0.1 in windows of 50 SNPs and steps of 5 SNPs). Ten independent unsupervised runs for K = 2 to 15 were performed, and for each value of K the run with maximum likelihood as estimated by the program is retained.

### EEMS

The EEMS analysis was conducted using all unrelated individuals, and SNPs were pruned for LD in the same way as in the PCA. As EEMS requires fine-scale geographically indexed samples, for the Sardinia EEMS analysis, we only used individuals whose four grandparents were all born in the same location at the town level. This resulted in 181 individuals across the island for analysis. For the Mediterranean region analysis, the merged dataset with Human Origins Array was used. Because of the scale of the Mediterranean region, we only used the two Sardinian populations of Cagliari and Sassari, the two Sardinian populations with the largest sample sizes that are geographically sufficiently distant to not be merged by EEMS. Populations from Human Origins Array data used in this analysis are: Spanish (Castilla y Leon, Castilla la Mancha, Extremadura, Cantabria, Cataluna, Valencia, Murcia, Andalucía, Baleares, Aragon, Galicia), Spanish_North, French_South, French, Bergamo, Italian_South, Tuscan, Sicilian, Mozabite, Algeria, Tunisian, and Spanish_Basque. We ran EEMS with different combinations of grid sizes and random seeds to assess robustness of results. We used the default settings for the EEMS hyper-parameters. For each run, we ran a burn-in of 1 million iterations followed by an additional 1 million iterations with posterior samples taken every thousandth iteration. We assessed the convergence of the MCMC chain by the posterior probability trace plot. We further assessed model fit by comparing the expected distance fitted by EEMS to the raw observed distances.

### f3/D

We restricted the f3 and D-statistic analyses to only a set of 156,237 SNPs ascertained in a single San individual as released by the Human Origin Array. For admixture f3 analyses, aimed to test for evidence of admixture in a target population, we computed the f3 statistic using all pairs of population from Europe (including Turkey/Greece, Italian peninsula and Iberian peninsula), Caucasus, Middle East, North Africa, and Subsaharan Africa (**Table S6**). For the outgroup f3 analyses, aimed to estimate the amount of shared drift between a pair of populations, we computed the f3 statistics between a Sardinian (Arzana or Cagliari) and another mainland population, while using the San individuals as the outgroup. Both f3 and D statistics were calculated using Admixtools version 3.0 (Patterson et al. 2012). Statistical significance were assessed using the default blocked jackknife implementation in Admixtools.

### ALDER

We also used an LD-based algorithm, ALDER, to test for admixture to complement the f3-based analyses. The full set of Human Origin Array SNPs were used, but with SNPs removed due to lack of recombination map information (based on sex-averaged deCODE map (Kong et al. 2010), or found in regions of long range LD (Price et al. 2008). For each Sardinian test populations, we tested all pairwise combinations of mainland populations as in the f3 admixture analysis.

In contrast to the approach taken by Loh et al., we opted to be conservative with interpreting ALDER results where the 2-reference LD decay fit is inconsistent with 1-reference LD decay fit from the program (*i.e*. a “successful fit” with warnings). We find that in general, successful fits where the 2-reference decay curve and the 1-reference decay curve agree with each other, the amplitude of the fit tend to be negatively correlated with the f3 statistics of the same source/target population triplets, even if the f3 statistics may be positive (*i.e*. suggesting no evidence of admixture). However, when the decay curve under the two scenarios do not agree with each other, the correlation with f3 statistics is also poorer and/or becomes positive. These results suggest complications from the shared past demography between source and target populations could have influenced the LD decay curve fitting (Loh et al. 2013). Thus we reported only successful fits for up to 5 pairs of populations with the highest estimated amplitude among those with significant evidence of admixture (P < 0.05 after multiple-testing correction by ALDER and Bonferroni correction for testing 15 Sardinian subpopulations), if available (**Table S4**). The pairs of source populations with the highest estimated amplitude of LD decay are the populations closest to the true ancestral populations (Loh et al. 2013) among those available in our analyses. We thus estimate the admixture proportion using these pairs of source population via f4 ratio test (Patterson et al. 2012) (**Table S4**).

### Estimating ancient population mixture proportions

For estimating mixture proportions in terms of a 3-way model of European admixture, we followed the method introduced in Haak et al (Haak et al. 2015). We estimated mixture proportion with respect to the early European farmers (LBK_EN), western hunter-gatherer (Loschbour), and the Yamnaya steppe herder (Yamnaya) published by Haak et al. The mixture proportion estimation was done using the command lsqlin in matlab, based on a matrix of relationships between the test sample, the three ancient reference samples and a set of 15 worldwide outgroups (Lazaridis et al. 2014, Haak et al. 2015). The 15 outgroups used are: Ami, Biaka, Bougainville, Chukchi, Eskimo, Han, Ju_hoan_North, Karitiana, Kharia, Mbuti, Onge, Papuan, She, Ulchi, and Yoruba. Our implementation was able to recapitulate the mixture proportions estimated in Haak et al. for the west Eurasian populations. We compared the fit using 2 reference samples (LBK_EN and Loschbour), and found that addition of Yamnaya ancestry marginally but significantly improved the fit (by extra sum of square F test, P-value < 10^−4^, with the exception of Gairo). Thus we present the 3-way mixture proportions in **Figure S10**.

### Identity-by-descent

We called IBD segments among the merged Sardinian-Human Origins dataset using RefinedIBD in Beagle 4, version r1274 (Browning and Browning 2013). Following the author’s recommendation, we set the ibdtrim parameter to 26 (corresponding to the number of markers per 0.15 cM), the overlap parameter to 260 (number of markers per 1.5cM). We also set the window parameter to 3386 (corresponding to sizes of 20cM). Three replicates with independent random starting seeds were generated, and merged using ibdmerge.jar utility provided by Beagle’s authors (https://faculty.washington.edu/browning/beagle_utilities/utilities.html). Visual inspections of IBD segments > 20Mb long showed that a lot of them overlapped regions of low marker densities. Thus we removed all segments overlapping 1Mb windows with marker density < 1x10^−5^ markers per bp, as well as segments with LOD scores < 6. This leaves a total of 5.63 million segments for analysis.

### MSMC

We used the multiple sequentially Markovian coalescent (msmc) to obtain historical effective population size trajectories and cross-coalescent rates through time. The msmc requires multiple deeply sequenced and phased haplotypes as input and outputs estimated coalescent rates in discretized time slices that can later be transformed to more interpretable values. We utilized a subset of unrelated individuals from Sardinia as well as a set of individuals from the 1000 Genomes project, sequenced by Complete Genomics (http://www.completegenomics.com/public-data/69-genomes/). These individuals are NA20502, NA20509, NA20510, and NA20511 from TSI, and NA12878, NA12891, NA12892, NA12877, NA12889, and NA12890 from CEU. For the 1000 genomes samples we called variants using the same pipeline that was applied to the 131 deeply sequenced Sardinians described above. Unphased variant callsets of each individual sample was phased with SHAPEIT2 using the 1000 Genomes Phase 3 reference panel (https://mathgen.stats.ox.ac.uk/impute/impute_v2.html#reference). Input files for msmc were then generated using custom scripts from the msmc github repository (https://github.com/stschiff/msmc-tools). As suggested by the authors of MSMC, we kept only regions where reads can be uniquely mapped (https://oc.gnz.mpg.de/owncloud/index.php/s/RNQAkHcNiXZz2fd). For effective population size inference, we used 4 individuals (8 phased haplotypes) from each population; for cross-coalescent rate inference we used pairs of two individuals from each population. We define the estimated divergence time between a pair of populations as first time point at which the cross-coalescent rate is at or above 0.5; for the range of the divergence time we used the first time point at which the cross-coalescent rate is at or above 0.25 and 0.75. The mutation rate used to scale time is 1.25x10^−8^ per bp per gen.

### Defining neutral regions of the genome for site-frequency spectrum analysis

We first removed all unsequenced sites and sites within 50kb of a coding exon or RNA genes in the human reference genome hg19. Of the remaining regions of the genome, we defined 5.5 million non-overlapping windows of size 100bp. We then computed the average recombination rate for each window (+/-10kb, using the deCODE recombination map (Kong et al. 2010). We opted to remove windows with exceptionally high or low recombination rates, keeping 826K windows with average recombination rate between 0.75 cM/Mb and 5 cM/Mb. We then removed all windows overlapping recombination hotspots (the deCODE recombination map at 10 kb resolution where the recombination rate is > 10 cM/Mb), or windows overlapping a conserved element (phastCons elements from 46 way vertebrate comparisons from UCSC genome browser). Of the remaining 763K windows, we assigned each window a percentile rank with respect to the average recombination rate, the B score measure of negative selection (McVicker et al. 2009), and by iHS (based on the weighted iHS signals from European, after remapping to hg19 coordinates using liftOver. Downloaded from http://hgdp.uchicago.edu/Browser_tracks/iHS/), and took the 10,000 windows with the lowest sum ranking. This produced a total of 10Mb sequence distributed across 3,039 continuous regions across the genome. Finally, we removed all CpG sites in these sequences, leaving 9,393,048 neutral sites for demographic inference using fastsimcoal.

### Demographic parameter estimation by Fastsimcoal

We estimated divergence time between Sardinia and mainland Europeans, as well as the growth rate since population split using a simulation-based framework implemented in Fastsimcoal (Excoffier et al. 2013). We constructed a 2-dimensional site frequency spectra among the ~9.4Mb neutral regions using deeply sequenced data from 67 Sardinians and 4 Tuscans from Complete Genomics. We downsampled the dataset to 100 Sardinian chromosomes and 8 Tuscan chromosomes, dropping remaining sites that did not have sufficient coverage. To avoid errors due to mis-annotation of the derived allele, we computed the 2-dimensional minor allele frequency spectra by using the minor allele based on the global frequency of the two alleles over all samples, per the convention established by fastsimcoal2. We then set up a demographic model, building on a previously published population size trajectory of Northern Europeans based on resequencing data (Gazave et al. 2014) (although we simplified the exponential growth phase reported in (Gazave et al. 2014) into 5 equal size epochs of constant population sizes equaling the harmonic mean of the exponentially growing size. Simulation showed this produced identical site frequency spectra). We assumed the Sardinians and the Tuscans diverged from the Northern European lineage at different times, each then started growing exponentially until the present day. The template file and the estimation file required by fastsimcoal are reproduced in the Supplemental Methods.

### Allele-sharing

We began by computing sharing at doubletons, as they comprise the rarest alleles that can be shared. We first collected a total of M variants that are seen exactly twice in the merged dataset of Sardinia and phase 3 of 1000 Genomes. Then for all pairs of populations *i* and *j*, each with relative fraction of sample sizes *x*_*i*_ and *x*_*j*_, we compute the ratio 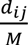, where *d*_*ij*_ is the number of doubletons shared between the two populations. This ratio is then normalized by 2*x*_*i*_*x*_*j*_ if *i ≠ j*, or 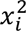if *i = j* to reflect the value of *d*_*ij*_ relative to what would be expected under random assignment of doubletons to populations, while accounting for uneven sample sizes.

For allele sharing at more frequent variants, we computed the allele-sharing ratios between pairs of populations as the probability that two randomly drawn carriers of the allele of a given minor allele frequency are from different populations, normalized by the panmictic expectation (Gravel et al. 2011, Nelson et al. 2012). Specifically, again define *x*_*i*_ and *x*_*j*_ as the relative fraction of sample sizes and *p*_*i*_ and *p*_*j*_ as the frequency of the minor allele in populations *i* and *j*. Then the probability of two randomly drawn carriers are from different populations is the probability of sampling two carriers from different populations over the total probability of sampling two carriers. In terms of the variables defined above, this is 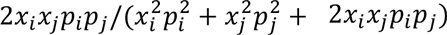. This quantity is normalized by the panmictic expectation, which is 2*x*_*i*_ *x*_*j*_.

We computed allele sharing statistics at two different levels of geographical groupings. At the province level, for each pair of populations with at least 70 unrelated individuals, we randomly selected 1 million variants and down-sampled the minor allele counts to 140 haploid chromosomes using a hypergeometric distribution, and then computed the allele-sharing ratio for 3 different minor allele frequency classes (0 to 0.01, 0.01 to 0.05, and 0.05 to 0.5). For analyses within Ogliastra, we computed the statistics for each pairs of populations with at least 17 unrelated individuals at three minor allele count classes (2, 3-5, and 6-34, corresponding to minor allele frequencies of 0.03, 0.075, and 0.5 out of 68 haploid chromosomes)

### Assessing sex-biased demography

We first used forward simulation on the basis of Sardinia population size history to evaluate the likelihood that the I2a1a1 haplotype drifted to current frequency of 39%. Given the population size trajectory estimated by MSMC using 4 unrelated high-coverage Sardinians from Arzana, we assumed an initial condition for the haplotype given a pair of parameters: the initial haplotype frequency, and the number of generations since its arrival in Sardinia. Then for all pairwise combinations of these two parameters we conducted forward simulation assuming the MSMC trajectory (with half of the Ne, as these are Y haplotypes), and in 100,000 simulations assessed the likelihood that a locus with the initializing condition would reach a present day frequency of 39%.

We then used two complementary approaches to assess possible sex-biased demography in the pre-history of Sardinia: (1) contrasting Sardinian-specific ancestry, and (2) contrasting the statistics D(Outgroup, Ancient, Sardinia or 1KG, CEU) computed on the X chromosome vs. the autosome.

For contrasting Sardinian-specific ancestry, we applied ADMIXTURE on similarly filtered SNP data from the autosome and X chromosome separately. Both sets of data were first filtered to retain only SNPs with minor allele frequencies > 0.02 in Europeans (1000 Genomes Europeans + 1,577 unrelated Sardinians). This left 6,740,788 SNPs on the autosome and 221,434 SNPs on the X chromosome. SNPs were then pruned by LD using 868 unrelated Sardinian females by removing one SNP of each pair of SNPs with r^2^ ≥0.1 (in windows of 50 SNPs and steps of 5 SNPs), leaving 433,704 autosomal SNPs and 18,918 X chromosome SNPs. We ran ADMIXTURE with K = 3, using all of unrelated Sardinians and TSI from 1000 Genomes. In general, TSI individuals form the first ancestry component, while the Sardinians are distributed in two different components as was observed in Figure 1B. We then compared between the autosome and X chromosome the distribution of the component showing the largest *F*_*St*_ from the TSI-dominated component to evaluate excess of the “Sardinian-specific” ancestry on X chromosome. In practice, this is equivalent to identifying the dominant ancestry component in isolated Sardinian groups such as Arzana. Significance was assessed by permuting individual ancestries 1 million times, as well as by bootstrapping individuals. Analysis was done on both male and females using Admixture v. 1.3 (Shringarpure et al. 2016), and confirmed by rerunning the analysis in the subset of 868 unrelated Sardinian females and in 839 non-Arzana Sardinian females. We also compared chr X to chr 7 only (**Figure S13**). SNPs were filtered similarly as above; in total we compared 26,164 SNPs on chr 7 to 18,918 SNPs on chr X.

For contrasting the D statistics on the X chromosome vs. the autosome, we first assembled a new merged dataset combining the Sardinians, 1000 genomes, and recently published high quality genomes of Neolithic farmers (Gamba et al. 2014, Lazaridis et al. 2014) (NE1 and Stuttgart) and Western hunter-gatherer (Lazaridis et al. 2014, Jones et al. 2015) (Bichon and Loschbour). To avoid ascertainment biases due to multi-sample genotype calling in our test populations, we evaluated the D-statistics across heterozygous sites found in 13 African individuals released by the Simons Genome Diversity Project [https://www.simonsfoundation.org/life-sciences/simons-genome-diversity-project-dataset/]. These 13 individuals were whole-genome sequenced to at least 30x, with minimal bias towards matching the human reference sequence, and were called on single-sample basis to avoid preferential calling of genotypes from populations with more individuals. Across the 13 individuals we ascertained 12.30M variants on the autosome, and 581.4K variants on the X chromosome. At each site, we sampled ten alleles (with replacement) from each of the ancient samples and constructed ten pseudoindividuals (i.e. based on the sampled allele we set a homozygous genotype at each position). We then merged the ancient samples with called genotypes from Sardinia and the 1000 Genomes for analysis. We report the mean across ten pseudoindividuals. Because the ascertained SNP sites were selected based on data generated using significantly different variant calling pipeline as used here, we opted to analyze only ascertained sites that are also polymorphic in both the Sardinian and 1000 Genomes dataset. We anticipate this approach to be conservative, as limiting the analysis to only polymorphic sites should bias the D statistics towards zero (*i.e*. inferring that Sardinia and CEU are more likely to be a clade). Indeed, our results comparing the X vs. A were not qualitatively changed if we assumed that everyone in the dataset are homozygous for the reference allele at uncalled ascertained sites. True or apparent triallelic SNPs were removed from analysis. Also, to avoid errors due to ancient DNA damage, we removed all C/T and A/G sites after merging. We then pruned the SNP set by LD using the same parameters as the Admixture analysis above. In total, 6.7K SNPs on the X chromosome and 182.9K SNPs on the autosome were used for analysis. We assessed the D statistic of the form of D(Outgroup, Ancient, Sardinia or 1KG, CEU) using admixtools v.3.0. Because admixtools could not properly handle hemizygous sites due to males in the dataset, the analysis was conducted using only females, so that chr X can be treated as an autosome. Individuals from 1000 Genomes YRI, LWK, ESN, PJL, and CHS were used in turn as outgroups. The ancient samples used were NE1, Stuttgart, Bichon, or Loschbour. Significance of the paired difference of the D statistics was assessed using 10,000 bootstrapping samples of the variant sites. We also repeated the analysis comparing chr X to chr 7 (N = 10,518 SNPs), observing no qualitative change to our results.

## SUPPLEMENTAL METHODS

Inference using fastsimcoal2 (version 2.5.11) was invoked with the following command:

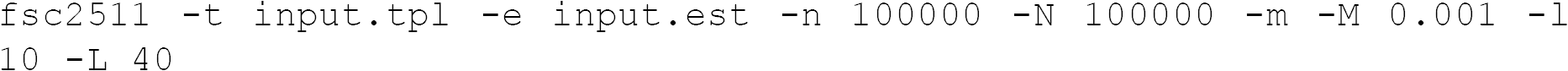

We report the estimated demographic parameter for the replicate with the maximum likelihood among 50 replicates of different starting seeds. To obtain confidence intervals of the estimated parameters we repeated the inference on 100 bootstrap replicates of the 2D spectra.

*Template file:*

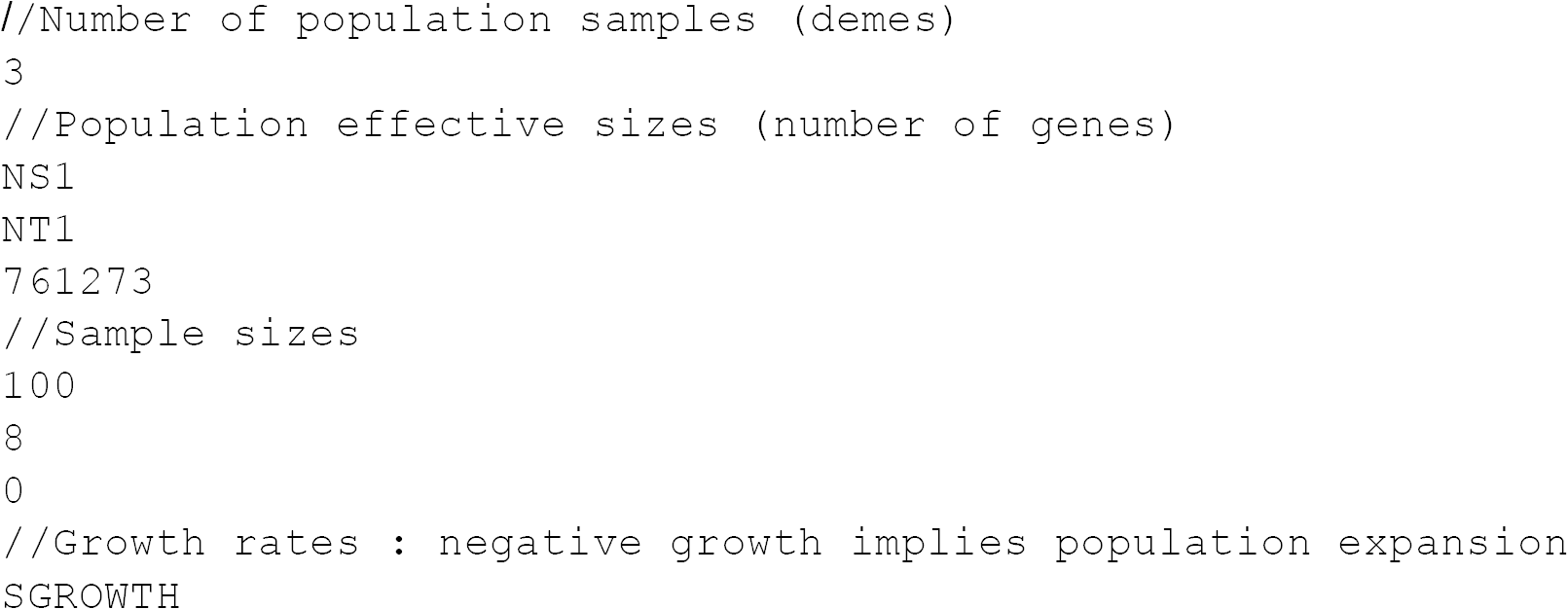

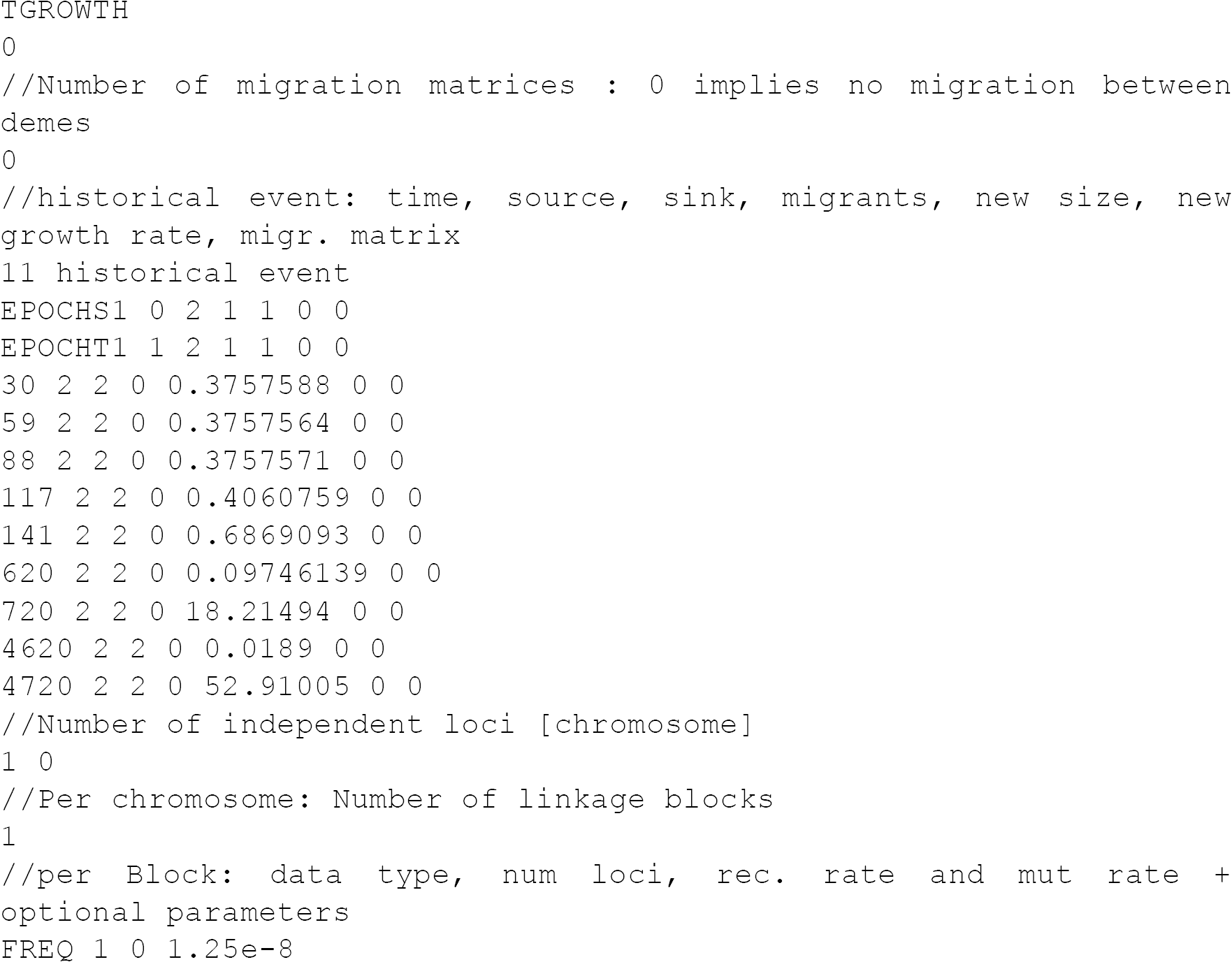

*Estimation file:*

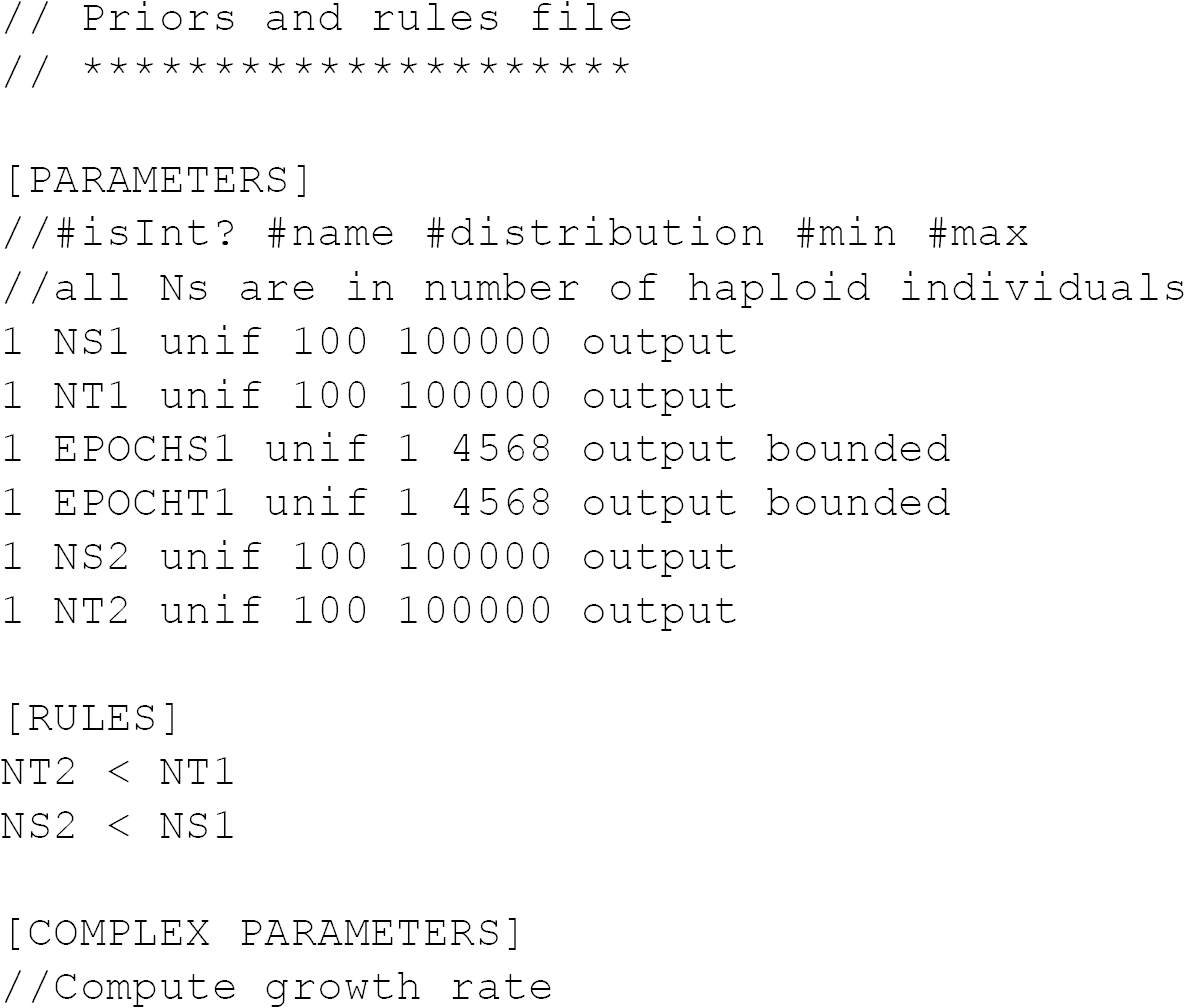

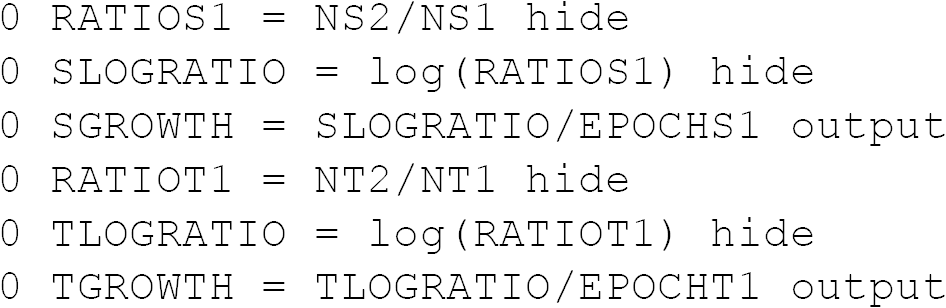

## REFERENCES CITED

Allentoft, M. E., M. Sikora, K. G. Sjogren, S. Rasmussen, M. Rasmussen, et al. (2015). “Population genomics of Bronze Age Eurasia.” Nature 522(7555): 167–172.

Barbujani, G., G. Bertorelle, G. Capitani and R. Scozzari (1995). “Geographical structuring in the mtDNA of Italians.” Proc Natl Acad Sci U S A 92(20): 9171–9175.

Barbujani, G. and R. R. Sokal (1990). “Zones of sharp genetic change in Europe are also linguistic boundaries.” Proc Natl Acad Sci U S A 87(5): 1816–1819.

Blasco Ferrer, E. (2010). Paleosardo: Le Radici Linguistiche Della Sardegna Neolítica, Walter De Gruyter Inc.

Botigue, L. R., B. M. Henn, S. Gravel, B. K. Maples, C. R. Gignoux, et al. (2013). “Gene flow from North Africa contributes to differential human genetic diversity in southern Europe.” Proc Natl Acad Sci U S A 110(29): 11791–11796.

Browning, B. L. and S. R. Browning (2009). “A unified approach to genotype imputation and haplotype-phase inference for large data sets of trios and unrelated individuals.” Am J Hum Genet 84(2): 210–223.

Browning, B. L. and S. R. Browning (2013). “Improving the accuracy and efficiency of identity-by-descent detection in population data.” Genetics 194(2): 459–471.

Bustamante, C. D. and S. Ramachandran (2009). “Evaluating signatures of sex-specific processes in the human genome.” Nat Genet 41(1): 8–10.

Calo, C. M., A. Melis, G. Vona and I. S. Piras (2008). “Sardinian Population (Italy): a Genetic Review.” International Journal of Modern Anthropology 1: 39–65.

Cann, H. M. (1998). “Human genome diversity.” C R Acad Sci III 321(6): 443–446.

Cao, A. and R. Galanello (2010). “Beta-thalassemia.” Genet Med 12(2): 61–76.

Caramelli, D., C. Vernesi, S. Sanna, L. Sampietro, M. Lari, et al. (2007). “Genetic variation in prehistoric Sardinia.” Hum Genet 122(3-4): 327–336.

Cavalli-Sforza, L. L. and A. Piazza (1993). “Human genomic diversity in Europe: a summary of recent research and prospects for the future.” Eur J Hum Genet 1(1): 3–18.

Chikhi, L., R. A. Nichols, G. Barbujani and M. A. Beaumont (2002). “Y genetic data support the Neolithic demic diffusion model.” Proc Natl Acad Sci U S A 99(17): 11008–11013.

Contu, D., L. Morelli, F. Santoni, J. W. Foster, P. Francalacci, et al. (2008). “Y-chromosome based evidence for pre-neolithic origin of the genetically homogeneous but diverse Sardinian population: inference for association scans.” PLoS One 3(1): e1430.

Dyson, S. L. and R. J. Rowland (2007). Archaeology and History in Sardinia from the Stone Age to the Middle Ages: Shepherds, Sailors, & Conquerors. Philadelphia, PA, University of Pennsylvania Museum of Archaeology and Anthropology.

Eaves, I. A., T. R. Merriman, R. A. Barber, S. Nutland, E. Tuomilehto-Wolf, et al. (2000). “The genetically isolated populations of Finland and sardinia may not be a panacea for linkage disequilibrium mapping of common disease genes.” Nat Genet 25(3): 320–323.

Excoffier, L., I. Dupanloup, E. Huerta-Sanchez, V. C. Sousa and M. Foll (2013). “Robust demographic inference from genomic and SNP data.” PLoS Genet 9(10): e1003905.

Francalacci, P., L. Morelli, A. Angius, R. Berutti, F. Reinier, et al. (2013). “Low-pass DNA sequencing of 1200 Sardinians reconstructs European Y-chromosome phylogeny.” Science 341(6145): 565–569.

Fraumene, C., E. Petretto, A. Angius and M. Pirastu (2003). “Striking differentiation of subpopulations within a genetically homogeneous isolate (Ogliastra) in Sardinia as revealed by mtDNA analysis.” Hum Genet 114(1): 1–10.

Gamba, C., E. R. Jones, M. D. Teasdale, R. L. McLaughlin, G. Gonzalez-Fortes, et al. (2014). “Genome flux and stasis in a five millennium transect of European prehistory.” Nat Commun 5: 5257.

Gazave, E., L. Ma, D. Chang, A. Coventry, F. Gao, et al. (2014). “Neutral genomic regions refine models of recent rapid human population growth.” Proc Natl Acad Sci U S A 111(2): 757762.

Ghirotto, S., S. Mona, A. Benazzo, F. Paparazzo, D. Caramelli, et al. (2010). “Inferring genealogical processes from patterns of Bronze-Age and modern DNA variation in Sardinia.” Mol Biol Evol 27(4): 875–886.

Goldberg, A., T. Gunther, N. A. Rosenberg and M. Jakobsson (2016). “Familial migration of the Neolithic contrasts massive male migration during Bronze Age in Europe inferred from ancient X chromosomes.” bioRxiv.

Gravel, S., B. M. Henn, R. N. Gutenkunst, A. R. Indap, G. T. Marth, et al. (2011). “Demographic history and rare allele sharing among human populations.” Proc Natl Acad Sci U S A 108(29): 11983–11988.

Gunther, T., C. Valdiosera, H. Malmstrom, I. Urena, R. Rodriguez-Varela, et al. (2015). “Ancient genomes link early farmers from Atapuerca in Spain to modern-day Basques.” Proc Natl Acad Sci U S A 112(38): 11917–11922.

Haak, W., I. Lazaridis, N. Patterson, N. Rohland, S. Mallick, et al. (2015). “Massive migration from the steppe was a source for Indo-European languages in Europe.” Nature 522(7555): 207211.

Hammer, M. F., F. L. Mendez, M. P. Cox, A. E. Woerner and J. D. Wall (2008). “Sex-biased evolutionary forces shape genomic patterns of human diversity.” PLoS Genet 4(9): e1000202.

Hellenthal, G., G. B. Busby, G. Band, J. F. Wilson, C. Capelli, et al. (2014). “A genetic atlas of human admixture history.” Science 343(6172): 747–751.

Henn, B. M., L. R. Botigue, S. Gravel, W. Wang, A. Brisbin, et al. (2012). “Genomic ancestry of North Africans supports back-to-Africa migrations.” PLoS Genet 8(1): e1002397.

Heyer, E., R. Chaix, S. Pavard and F. Austerlitz (2012). “Sex-specific demographic behaviours that shape human genomic variation.” Mol Ecol 21(3): 597–612.

Hofmanova, Z., S. Kreutzer, G. Hellenthal, C. Sell, Y. Diekmann, et al. (2016). “Early farmers from across Europe directly descended from Neolithic Aegeans.” Proc Natl Acad Sci U S A 113(25): 6886–6891.

International HapMap, C., D. M. Altshuler, R. A. Gibbs, L. Peltonen, D. M. Altshuler, et al. (2010). “Integrating common and rare genetic variation in diverse human populations.” Nature 467(7311): 52–58.

Jones, E. R., G. Gonzalez-Fortes, S. Connell, V. Siska, A. Eriksson, et al. (2015). “Upper Palaeolithic genomes reveal deep roots of modern Eurasians.” Nat Commun 6: 8912.

Joshi, P. K., T. Esko, H. Mattsson, N. Eklund, I. Gandin, et al. (2015). “Directional dominance on stature and cognition in diverse human populations.” Nature 523(7561): 459–462.

Keinan, A. and A. G. Clark (2012). “Recent explosive human population growth has resulted in an excess of rare genetic variants.” Science 336(6082): 740–743.

Keinan, A., J. C. Mullikin, N. Patterson and D. Reich (2009). “Accelerated genetic drift on chromosome X during the human dispersal out of Africa.” Nat Genet 41(1): 66–70.

Keller, A., A. Graefen, M. Ball, M. Matzas, V. Boisguerin, et al. (2012). “New insights into the Tyrolean Iceman’s origin and phenotype as inferred by whole-genome sequencing.” Nat Commun 3: 698.

Kong, A., G. Thorleifsson, D. F. Gudbjartsson, G. Masson, A. Sigurdsson, et al. (2010). “Fine-scale recombination rate differences between sexes, populations and individuals.” Nature 467(7319): 1099–1103.

Lazaridis, I., N. Patterson, A. Mittnik, G. Renaud, S. Mallick, et al. (2014). “Ancient human genomes suggest three ancestral populations for present-day Europeans.” Nature 513(7518): 409–413.

Lettre, G. and J. N. Hirschhorn (2015). “Small island, big genetic discoveries.” Nat Genet 47(11): 1224–1225.

Li, J. Z., D. M. Absher, H. Tang, A. M. Southwick, A. M. Casto, et al. (2008). “Worldwide human relationships inferred from genome-wide patterns of variation.” Science 319(5866): 1100–1104.

Loh, P. R., M. Lipson, N. Patterson, P. Moorjani, J. K. Pickrell, et al. (2013). “Inferring admixture histories of human populations using linkage disequilibrium.” Genetics 193(4): 1233–1254.

Lohmueller, K. E. (2014). “The impact of population demography and selection on the genetic architecture of complex traits.” PLoS Genet 10(5): e1004379.

Marrosu, M. G., C. Motzo, R. Murru, R. Lampis, G. Costa, et al. (2004). “The co-inheritance of type 1 diabetes and multiple sclerosis in Sardinia cannot be explained by genotype variation in the HLA region alone.” Hum Mol Genet 13(23): 2919–2924.

Mathieson, I., I. Lazaridis, N. Rohland, S. Mallick, N. Patterson, et al. (2015). “Genome-wide patterns of selection in 230 ancient Eurasians.” Nature 528(7583): 499–503.

McVicker, G., D. Gordon, C. Davis and P. Green (2009). “Widespread genomic signatures of natural selection in hominid evolution.” PLoS Genet 5(5): e1000471.

Moorjani, P., N. Patterson, J. N. Hirschhorn, A. Keinan, L. Hao, et al. (2011). “The history of African gene flow into Southern Europeans, Levantines, and Jews.” PLoS Genet 7(4): e1001373.

Morelli, L., D. Contu, F. Santoni, M. B. Whalen, P. Francalacci, et al. (2010). “A comparison of Y-chromosome variation in Sardinia and Anatolia is more consistent with cultural rather than demic diffusion of agriculture.” PLoS One 5(4): e10419.

Morelli, L., M. G. Grosso, G. Vona, L. Varesi, A. Torroni, et al. (2000). “Frequency distribution of mitochondrial DNA haplogroups in Corsica and Sardinia.” Hum Biol 72(4): 585–595.

Naitza, S., E. Porcu, M. Steri, D. D. Taub, A. Mulas, et al. (2012). “A genome-wide association scan on the levels of markers of inflammation in Sardinians reveals associations that underpin its complex regulation.” PLoS Genet 8(1): e1002480.

Nelson, M. R., D. Wegmann, M. G. Ehm, D. Kessner, P. St Jean, et al. (2012). “An abundance of rare functional variants in 202 drug target genes sequenced in 14,002 people.” Science 337(6090): 100–104.

Novembre, J. and M. Stephens (2008). “Interpreting principal component analyses of spatial population genetic variation.” Nat Genet 40(5): 646–649.

Olivieri, A., C. Sidore, A. Achilli and A. Angius (2017). “Mitogenomes of Both Western European and Near Eastern Ancestry Were Present in Pre-Neolithic Sardinia: Implications for the Genetic Origin of Modern Europeans.” unpublished.

Pala, M., A. Achilli, A. Olivieri, B. Hooshiar Kashani, U. A. Perego, et al. (2009). “Mitochondrial haplogroup U5b3: a distant echo of the epipaleolithic in Italy and the legacy of the early Sardinians.” Am J Hum Genet 84(6): 814–821.

Paschou, P., P. Drineas, E. Yannaki, A. Razou, K. Kanaki, et al. (2014). “Maritime route of colonization of Europe.” Proc Natl Acad Sci U S A 111(25): 9211–9216.

Passarino, G., P. A. Underhill, L. L. Cavalli-Sforza, O. Semino, G. M. Pes, et al. (2001). “Y chromosome binary markers to study the high prevalence of males in Sardinian centenarians and the genetic structure of the Sardinian population.” Hum Hered 52(3): 136–139.

Patterson, N., P. Moorjani, Y. Luo, S. Mallick, N. Rohland, et al. (2012). “Ancient admixture in human history.” Genetics 192(3): 1065–1093.

Petkova, D., J. Novembre and M. Stephens (2016). “Visualizing spatial population structure with estimated effective migration surfaces.” Nat Genet 48(1): 94–100.

Pilia, G., W. M. Chen, A. Scuteri, M. Orru, G. Albai, et al. (2006). “Heritability of cardiovascular and personality traits in 6,148 Sardinians.” PLoS Genet 2(8): e132.

Pistis, G., I. Piras, N. Pirastu, I. Persico, A. Sassu, et al. (2009). “High differentiation among eight villages in a secluded area of Sardinia revealed by genome-wide high density SNPs analysis.” PLoS One 4(2): e4654.

Price, A. L., M. E. Weale, N. Patterson, S. R. Myers, A. C. Need, et al. (2008). “Long-range LD can confound genome scans in admixed populations.” Am J Hum Genet 83(1): 132–135; author reply 135-139.

Pugliatti, M., G. Rosati, H. Carton, T. Riise, J. Drulovic, et al. (2006). “The epidemiology of multiple sclerosis in Europe.” Eur J Neurol 13(7): 700–722.

Raghavan, M., P. Skoglund, K. E. Graf, M. Metspalu, A. Albrechtsen, et al. (2014). “Upper Palaeolithic Siberian genome reveals dual ancestry of Native Americans.” Nature 505(7481): 87–91.

Rootsi, S., C. Magri, T. Kivisild, G. Benuzzi, H. Help, et al. (2004). “Phylogeography of Y-chromosome haplogroup I reveals distinct domains of prehistoric gene flow in europe.” Am J Hum Genet 75(1): 128–137.

Sanna, S., M. Pitzalis, M. Zoledziewska, I. Zara, C. Sidore, et al. (2010). “Variants within the immunoregulatory CBLB gene are associated with multiple sclerosis.” Nat Genet 42(6): 495-497.

Scherer, S. (2010). Guide to the Human Genome, Cold Spring Harbor Laboratory Press.

Schiffels, S. and R. Durbin (2014). “Inferring human population size and separation history from multiple genome sequences.” Nat Genet 46(8): 919–925.

Schraiber, J. G. and J. M. Akey (2015). “Methods and models for unravelling human evolutionary history.” Nat Rev Genet 16(12): 727–740.

Semino, O., G. Passarino, P. J. Oefner, A. A. Lin, S. Arbuzova, et al. (2000). “The genetic legacy of Paleolithic Homo sapiens sapiens in extant Europeans: a Y chromosome perspective.” Science 290(5494): 1155–1159.

Shringarpure, S. S., C. D. Bustamante, K. Lange and D. H. Alexander (2016). “Efficient analysis of large datasets and sex bias with ADMIXTURE.” BMC Bioinformatics 17: 218.

Sidore, C., F. Busonero, A. Maschio, E. Porcu, S. Naitza, et al. (2015). “Genome sequencing elucidates Sardinian genetic architecture and augments association analyses for lipid and blood inflammatory markers.” Nat Genet 47(11): 1272–1281.

Sikora, M., M. L. Carpenter, A. Moreno-Estrada, B. M. Henn, P. A. Underhill, et al. (2014). “Population genomic analysis of ancient and modern genomes yields new insights into the genetic ancestry of the Tyrolean Iceman and the genetic structure of Europe.” PLoS Genet 10(5): e1004353.

Simons, Y. B., M. C. Turchin, J. K. Pritchard and G. Sella (2014). “The deleterious mutation load is insensitive to recent population history.” Nat Genet 46(3): 220–224.

Skoglund, P., H. Malmstrom, M. Raghavan, J. Stora, P. Hall, et al. (2012). “Origins and genetic legacy of Neolithic farmers and hunter-gatherers in Europe.” Science 336(6080): 466–469.

Uricchio, L. H., N. A. Zaitlen, C. J. Ye, J. S. Witte and R. D. Hernandez (2016). “Selection and explosive growth alter genetic architecture and hamper the detection of causal rare variants.” Genome Res 26(7): 863–873.

Vona, G. (1997). “The peopling of Sardinia (Italy): history and effects.” International Journal of Anthropology 12(1): 71–87.

Wilkins, J. F. and F. W. Marlowe (2006). “Sex-biased migration in humans: what should we expect from genetic data?” Bioessays 28(3): 290–300.

Zavattari, P., E. Deidda, M. Whalen, R. Lampis, A. Mulargia, et al. (2000). “Major factors influencing linkage disequilibrium by analysis of different chromosome regions in distinct populations: demography, chromosome recombination frequency and selection.” Hum Mol Genet 9(20): 2947–2957.

Zei, G., A. Lisa, O. Fiorani, C. Magri, L. Quintana-Murci, et al. (2003). “From surnames to the history of Y chromosomes: the Sardinian population as a paradigm.” Eur J Hum Genet 11(10): 802–807.

Zoledziewska, M., G. Costa, M. Pitzalis, E. Cocco, C. Melis, et al. (2009). “Variation within the CLEC16A gene shows consistent disease association with both multiple sclerosis and type 1 diabetes in Sardinia.” Genes Immun 10(1): 15–17.

Zoledziewska, M., C. Sidore, C. W. Chiang, S. Sanna, A. Mulas, et al. (2015). “Height-reducing variants and selection for short stature in Sardinia.” Nat Genet 47(11): 1352–1356.

